# Meiotic nuclear architecture in distinct mole vole hybrids with Robertsonian translocations: chromosome chains, stretched centromeres, and distorted recombination

**DOI:** 10.1101/2020.09.17.301473

**Authors:** Sergey Matveevsky, Artemii Tretiakov, Irina Bakloushinskaya, Anna Kashintsova, Oxana Kolomiets

## Abstract

Genome functioning in hybrids faces inconsistency. This mismatch is manifested clearly in meiosis during chromosome synapsis and recombination. Species with chromosomal variability can be a model for exploring genomic battles with high visibility due to the use of advanced immunocytochemical methods. We studied synaptonemal complexes (SC) and prophase I processes in 44-chromosome intraspecific (*Ellobius tancrei* × *E. tancrei*) and interspecific (*Ellobius talpinus* × *E. tancrei*) hybrid mole voles heterozygous for 10 Robertsonian translocations. The same pachytene failures were found for both types of hybrids. In the intraspecific hybrid, the chains were visible in the pachytene stage, then 10 closed SC trivalents formed in the late pachytene and diplotene stage. In the interspecific hybrid, as a rule, SC trivalents composed the SC chains and rarely could form closed configurations. Metacentrics involved with SC trivalents had stretched centromeres in interspecific hybrids. Linkage between neighboring SC trivalents was maintained by stretched centromeric regions of acrocentrics. This centromeric plasticity in structure and dynamics of SC trivalents was found for the first time. We assume that stretched centromeres were a marker of altered nuclear architecture in heterozygotes due to differences in the ancestral chromosomal territories of the parental species. Restructuring of the intranuclear organization and meiotic disturbances can contribute to the sterility of interspecific hybrids, and lead to the reproductive isolation of studied species.

**Author summary:** Meiosis is essential for sexual reproduction to produce haploid gametes. Prophase I represents a crucial meiotic stage because key processes such as chromosomal pairing, synapsis and desynapsis, recombination, and transcriptional silencing occur at this time. Alterations in each of these processes can activate meiotic checkpoints and lead to the elimination of meiocytes. Here we have shown that two groups of experimental hybrids, intraspecific and interspecific—which were heterozygous for 10 identical Robertsonian translocations—had pachytene irregularities and reduced recombination. However, intraspecific and interspecific hybrids exhibited different patterns of synaptonemal complex (SC) trivalent behavior. In the former, open SC trivalents comprised SC chains due to heterosynapsis of short arms of acrocentrics in early and mid-pachytene and were then able to form 2–4 and even 7 and 10 closed SC trivalents in the late pachytene and diplotene stages. In the second mole voles, SC trivalents had stretched centromeres of the metacentrics, and chains of SC trivalents were formed due to stretched centromeres of acrocentrics. Such compounds could not lead to the formation of separate closed SC trivalents. The distant ancestral points of chromosome attachment with a nuclear envelope in the heterozygous nuclei probably lead to stretching of SC trivalents and their centromeric regions, which can be regarded as an indicator of the reorganization of the intranuclear chromatin landscape. These abnormalities, which were revealed in in prophase I, contribute to a decrease the fertility of intraspecific mole voles and promote the sterility of interspecific mole voles.

## Introduction

Genome integrity is crucial for a species; its specificity is supported by reproductive isolation. Hybrid sterility may develop between species or genetically differentiated populations, as a primary or secondary feature of reproductive isolation (Dobzhansky, 1937, Benirschke, 1967, Basrur, 1969, White, 1977, Pavlova and Searle, 2018). Two genetic materials in heterozygous admixed genome interact in various compositions (Runemark et al., 2019) and these hybrid states are often referred to as “genomic conflict” (Johnson, 2010, Crespi and Nosil, 2013), “genomic shock” (McClintock, 1984, Ha et al., 2009), “genomic stress” (McClintock 1984, Petrov et al., 1995), or “nucleus at war” (Jones and Langdon, 2013).

Hybrid incompatibility (Watkins, 1932, Dobzhansky, 1937, Muller, 1942) is expressed either by visible or cryptic changes in the phenotype (Hale et al., 1993, Foreit, 1996, Ishishita et al., 2015) or disturbances in chromosome sets and gene expression and alterations in the gene networks (Haerty and Singh, 2006, Landry et al., 2007). Epistatic interactions between chromosomal regions (heterochromatin blocks, centromeres, and telomeres), satellite DNA, small RNA, and epigenetic chromatin modifications (Brown, O’Neill, 2010) are essential for the emergence of Dobzhansky–Muller incompatibility. Hybrid and heterozygous animals are excellent models for studying the effect of chromosomal rearrangements on the development of the organism, cellular function, intracellular structures, germ cell formation— including the most important stage, meiosis—and, in general, chromosomal evolution. Chromosome differences in hybrid meiocytes can manifest as various irregularities in chromosome synapsis, recombination, chromatin landscape, and transcriptional inactivation (Johannisson and Winking, 1994, 1998, Sharma et al., 2003, Bhattacharyya et al., 2013).

Chromosome heterozygosity can be studied either in laboratory hybrids (Forejt et al., 1980, Gill, 1980, Matsuda et al., 1992, Matveevsky et al., 2014, Gureeva et al., 2016) or in natural hybrids, for example, from hybrid zones (Ivanitskaya et al., 2010, Matveevsky et al., 2012, Lin et al., 2018). Experimental hybridization allows researchers to obtain offspring from parental forms with known karyotypic data, which facilitates a more accurate assessment of the effect of structural hybridity on the processes of cell division, morphology, and fertility (Chang et al., 1969, King, 1993).

Mole voles are gratifying sources for experimental hybridization. One of the three cryptic species has a stable karyotype, namely *Ellobius talpinus* (2n = 54, NF = 54), and no natural hybrids are known (Lebedev et al., 2020). Two other species have Robertsonian (Rb) chromosome variability: *Ellobius tancrei* (2n = 54–30, NF = 56) and *Ellobius alaicus* (2n = 52–48, NF = 56) (Vorontsov et al., 1969; Lyapunova et al., 1984, 2010, Bakloushinskaya et al., 2013, 2019). These species demonstrate intra- and interspecific hybridization (Bakloushinskaya et al., 2019, Romanenko et al., 2019). The 54-chromosome karyotypes of *E. talpinus* and *E. tancrei* are identical and differ only in chromosome #7 due to centromere repositioning (Romanenko et al., 2007). Ten pairs of bi-armed (Rb) chromosomes are a feature of the 34-chromosome karyotype of *E. tancrei* (Romanenko et al., 2019) (Fig 1). Karyotypic variability allows researchers to simulate different natural chromosomal combinations in experimental hybrids (Kolomiets et al., 1983, 1985, 1986, Lyapunova and Yakimenko, 1985, Lyapunova et al., 1990, Bakloushinskaya et al., 2010, 2012, Matveevsky et al., 2015, 2017). The first description of chromosome chains was made for an intraspecific *E. tancrei* hybrid heterozygous for 10 Rb translocations (Kolomiets et al., 1985, Bogdanov et al., 1986). It would be extremely interesting to compare the heterozygotes with the same diploid numbers obtained from crossing *E. tancrei* chromosomal forms between themselves and between two species, *E. talpinus* and *E. tancrei*.

**Fig 1.**
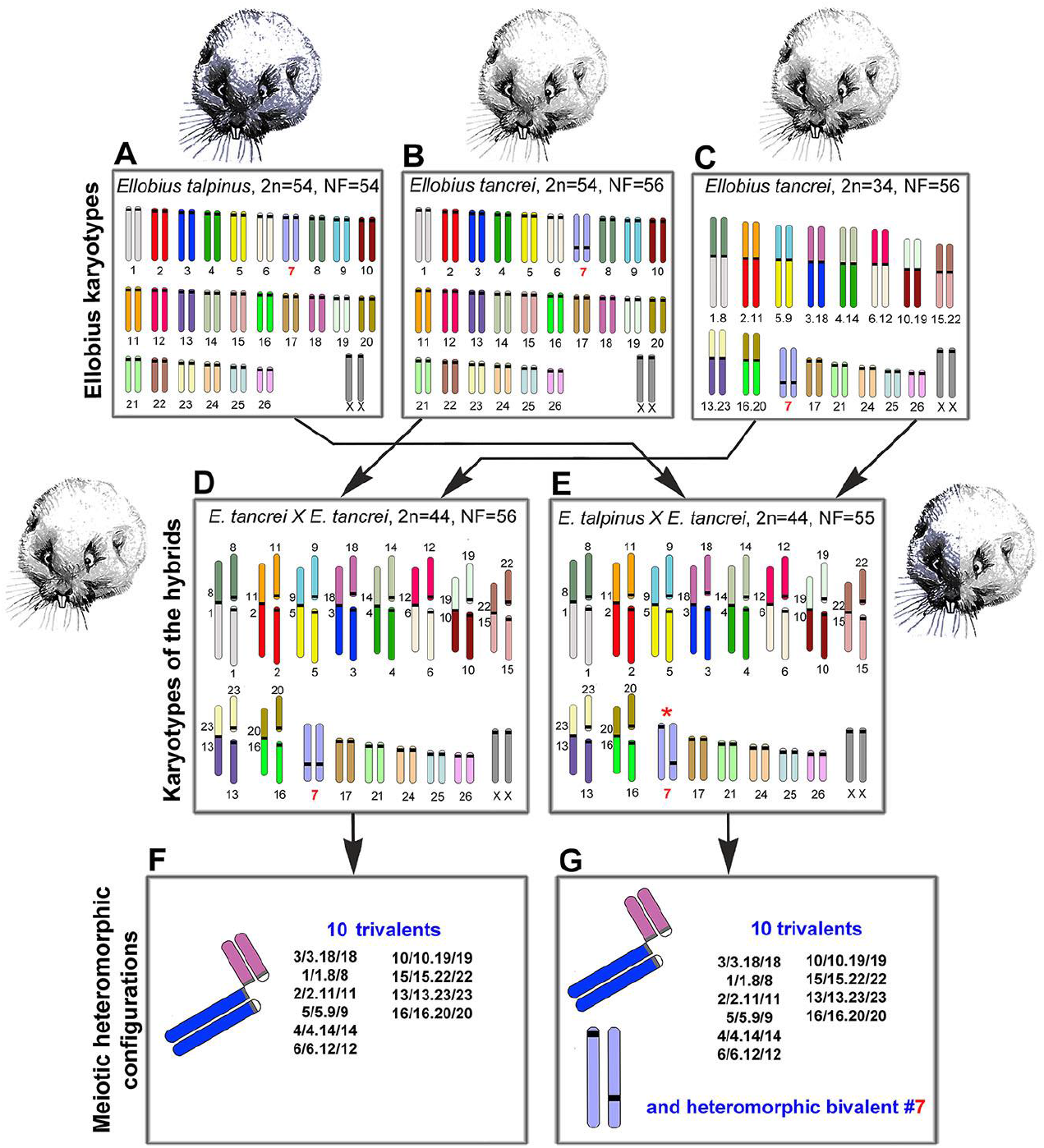
Scheme of the experimental hybridization of *Ellobius* species and hybrids. **(A–C)** Karyotypes for *E talpinus*, 2n = 54, NF = 54 (A); *E. tancrei*, 2n = 54, NF = 56 (B); and *E. tancrei*, 2n = 34, NF = 56 (C). **(D**, **E)** F1 hybrid karyotypes for *E. tancrei* × *E. tancrei*, 2n = 44, NF = 56 (D) and *E. talpinus* × *E. tancrei*, 2n = 44, NF = 55 (E). **(F**, **G)** Chromosome heteromorphic configurations in meiotic prophase I of F1 hybrids: 10 trivalents (F), and 10 trivalents and the heteromorphic chromosome #7 (G).

In this study, we first compared chromosome synapsis and recombination in intra- and interspecific hybrids with the same chromosome number and with numerous translocations. We analyzed 44-chromosome F1 hybrids, intraspecific *E. tancrei* × *E. tancrei*, and interspecific *E. talpinus* × *E. tancrei*, heterozygous for 10 identical Rb translocations (Fig 1, compare D and E). Each of the hybrids is expected to form 10 identical SC trivalents. A comparative analysis of chromosome synapsis, recombination, and meiotic silencing in the fertile and sterile hybrids allowed us to assess the effect of chromosome rearrangements on these processes, as well as to draw conclusions about the degree of species divergence.

## Results

### Experimental hybridization

We studied parental mole vole species—*E. talpinus* (2n = 54, NF = 54), *E. tancrei* (2n = 54, NF = 56), and another form of *E. tancrei* (2n = 34, NF = 56)—and F1 hybrids—intraspecific *E. tancrei* × *E. tancrei* (2n = 44, NF = 56) and interspecific *E. talpinus* × *E. tancrei* (2n = 44, NF = 55) (Fig 1).

The karyotypes of interspecific and intraspecific hybrids differ little from each other. They include 5 pairs of acrocentrics, 10 metacentrics and 20 acrocentrics, which are homologous to the arms of metacentrics, pair #7 and a pair of isomorphic sex (XX) chromosomes. Two types of hybrids differ only by the chromosome #7 pair. In the interspecific hybrid, this pair is heteromorphic (Fig 1), while in the intraspecific hybrid, it is represented by a pair of homologous submetacentrics. In total, we detected 10 SC trivalents, 6 SC bivalents, including a heteromorphic SC bivalent in the interspecific hybrid, and a sex XX bivalent in meiotic prophase I in spermatocyte spreads and squashes of both hybrids (Fig 1).

The gonadosomatic index (GSI) can be used as an indicator of the state of the reproductive system. It is calculated as the ratio of the weight of the testes to the weight of the body (Adebayo et al., 2009). This parameter is species specific; varies depending on age, stage of development, sex, and breeding season (Pochron et al., 2002); and may reflect the rates of sperm production as well as sperm function (Gomendio et al., 2006). Comparison of the parameter in closely related forms, species, and hybrids can be informative. Hence, we calculated the GSIs in adult mole voles, parental species, and hybrids. GSI (mean [M] ± standard deviation [SD]) in the interspecific hybrid (0.11 ± 0.04, n = 3) was approximately half of the *E. talpinus* GSI (0.20 ± 0.02, n = 3) and approximately one third of the *E. tancrei* (2n = 34) GSI (0.31 ± 0.10, n = 3). Species and hybrids significantly differ from each other in this parameter (p < 0.05). This indicator suggests that interspecific hybrids have reproductive dysfunction. Unfortunately, we do not have data on GSI of intraspecific hybrids.

There were numerous spermatocytes and mature spermatids in the testicular cell suspension of the parental species and the intraspecific hybrids (Fig S1A, S1B, S1C, and S1D). In the interspecific hybrids, the number of spermatocytes was significantly lower, and mature spermatids and spermatozoa were not detected at all, which was confirmed by histological examination of testicular tissue sections (Fig S1B, S1C, and S1D). Testicles of the interspecific hybrids were significantly smaller than in 34-chromosome parental forms (Fig S1E). Testis weights (and most likely testis volumes) < 55% of normal values indicate sterility (de Boer and de Jong, 1989). None of the 93 interspecific F1 *E. talpinus* × *E. tancrei* hybrids produced offspring (shown graphically in Fig S1F, compare all species and hybrids). The fertility data of mole voles in this work are in good agreement with the previous data (Lyapunova and Yakimenko, 1985). Moreover, for the intraspecific hybrids, there was an increased time interval from the moment of coupling to the first litter and an increased period between births compared with the parental forms (Lyapunova and Yakimenko, 1985). Thus, the intraspecific hybrid had slightly reduced fertility, and the interspecific hybrid was sterile.

### Synaptic behavior of chromosomes

We first described the chromosome behavior in pachytene spermatocytes of parental forms. In *E. talpinus*, there were 26 acrocentric bivalents and a sex (XX) bivalent, covered by a γH2AFX cloud, that moved to the periphery of the nucleus (Fig 2A and S2A, S2B, and S2C). In 54-chromosome *E. tancrei*, there were 25 acrocentric SC bivalents and 1 submetacentric SC bivalent, and a XX bivalent, covered by a γH2AFX cloud, that moved to the periphery of the nucleus (Fig 2B S2D, S2E, and S2F). In 34-chromosome *E. tancrei*, there were 10 metacentric SC bivalents, 1 submetacentric SC bivalent #7, and 5 acrocentric bivalents, and a XX bivalent, covered by a γH2AFX cloud, that moved to the nuclear periphery (Fig 2C S2G, S2H, and S2I).

**Fig 2.**
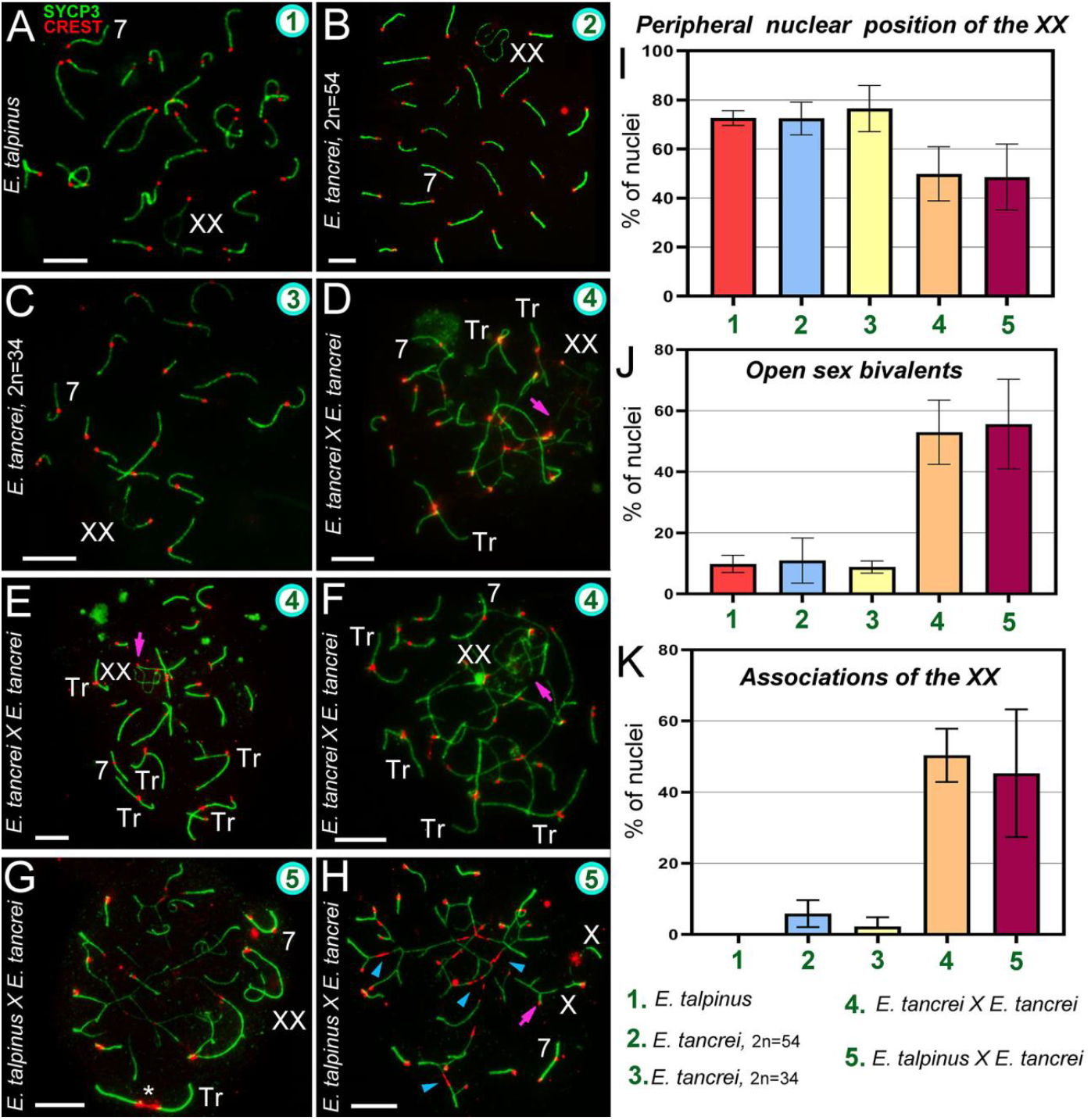
Chromosome synapsis and irregularities in pachytene spermatocytes of *Ellobius* species and hybrids. The green numbers in the micrographs and diagrams correspond to parental species and hybrids (see captions under diagram K). Chromosome #7 was clearly identified in all nuclei (A–H). **(A)** *Ellobius talpinus:* 26 synaptonemal complexes (SCs) and 1 XX bivalent. **(B)** *E. tancrei* (2n = 54): 26 SCs and 1 XX bivalent. **(C)** *E. tancrei* (2n = 34): 16 SCs and 1 XX bivalent. **(D–F)** Intraspecific hybrid *E. tancrei* × *E. tancrei* (2n = 44): No nucleus had the expected chromosome formula (6 SCs, 1 XX bivalent, 10 SC trivalents). There were several SCs, 3–5 free SC trivalents, a XX bivalent, and SC trivalents in chains. **(G, H)** Interspecific hybrid *E. talpinus* × *E. tancrei* (2n = 44): Expected chromosome formula: 6 SCs, 1 XX bivalent, 10 SC trivalents. One nucleus contained 3 SCs, 1 XX bivalent, 1 SC trivalent with a stretched centromere (white star), and SC trivalents in a chain (G). The other nucleus had 6 SCs, 1 XX bivalent, and SC trivalents in a chain (no free SC trivalents) (H). Some metacentrics in the SC trivalents had a stretched centromere (blue arrowheads). **(I)** The number of cells in which the XX bivalent moved to the periphery of the nuclei: Hybrids had significantly lower rates compared with the parent species (see Table S1). **(J)** The number of cells with an open XX bivalent: This parameter was much higher in hybrids than in parents (Table S1 indicates significant differences). This parameter increased from intraspecific hybrids to interspecific hybrids (differences between them were not significant). For examples of open XX bivalent, see D, E, and H. **(K)** The number of cells with associations of XX with autosomes and trivalents: There were significant differences between parents and hybrids, and this parameter was slightly higher in the intraspecific hybrid (2n = 44) than in the other hybrid (see Table S1). An open sex bivalent associated with trivalents are shown in D, E, and H. Scale bars represent 5 μm.

In an intraspecific hybrid, there were 10 closed and open SC trivalents, 5 acrocentric bivalents, 1 submetacentric bivalent #7, and a XX bivalent (Fig 2D, 2E and 2F S3A-, S3B, and S3C). The γH2AFX cloud completely covered the XX bivalent and asynaptic regions of the open SC trivalent (Fig S2J, S2K, and S2L). In the interspecific hybrid, there were 10 closed and open SC trivalents, 5 acrocentric bivalents, 1 heteromorphic chromosome #7 bivalent, and a XX bivalent (Fig 2G and 2H and S4A). The γH2AFX cloud completely covered the XX bivalent and some asynaptic regions of open SC trivalents (Fig S2M, S2N, and S2O).

The XX bivalent, as a rule, had two telomeric synaptic sites and a large central asynaptic area (closed configuration). A chromatin-dense body (ChB) was formed in one of the axial elements of the XX’ extended area of asynapsis (Fig S3D and S4C) (Kolomiets et al., 1991, Matveevsky et al., 2016). One of the synaptic sites may have been absent when the axis or axes were associated with SC trivalents or bivalents (open configuration of XX bivalent).

In mammals, sex bivalents are usually moved to the periphery of meiotic nuclei and covered in a cloud of protein inactivators of asynaptic chromatin. In hybrids, the XX bivalent was significantly less likely to move to the nucleus periphery than in the parents. However, the difference between intra- and interspecific hybrids was not statistically significant (Fig 2I, Table S1[2]). The number of open XX bivalents in hybrids was higher than in parents; the difference between parents and hybrids was significant, but not significant within each of the groups (Fig 2E, 2H, and 2J, S1(3) Table, S3C, S3E and S4F).

### Synapsis defects and associations of the XX and SC trivalents in hybrids

Chromosome synapsis defects included atypical lengthening of the short arms of acrocentrics of the SC trivalent in both hybrids (Fig 3H, S3B, and S4H), as well as atypical lengthening of one of the synaptic regions of the sex bivalent in the intraspecific hybrid (Fig 3H). This lengthening occurred due to a slight stretch at the centromeric region.

**Fig 3.**
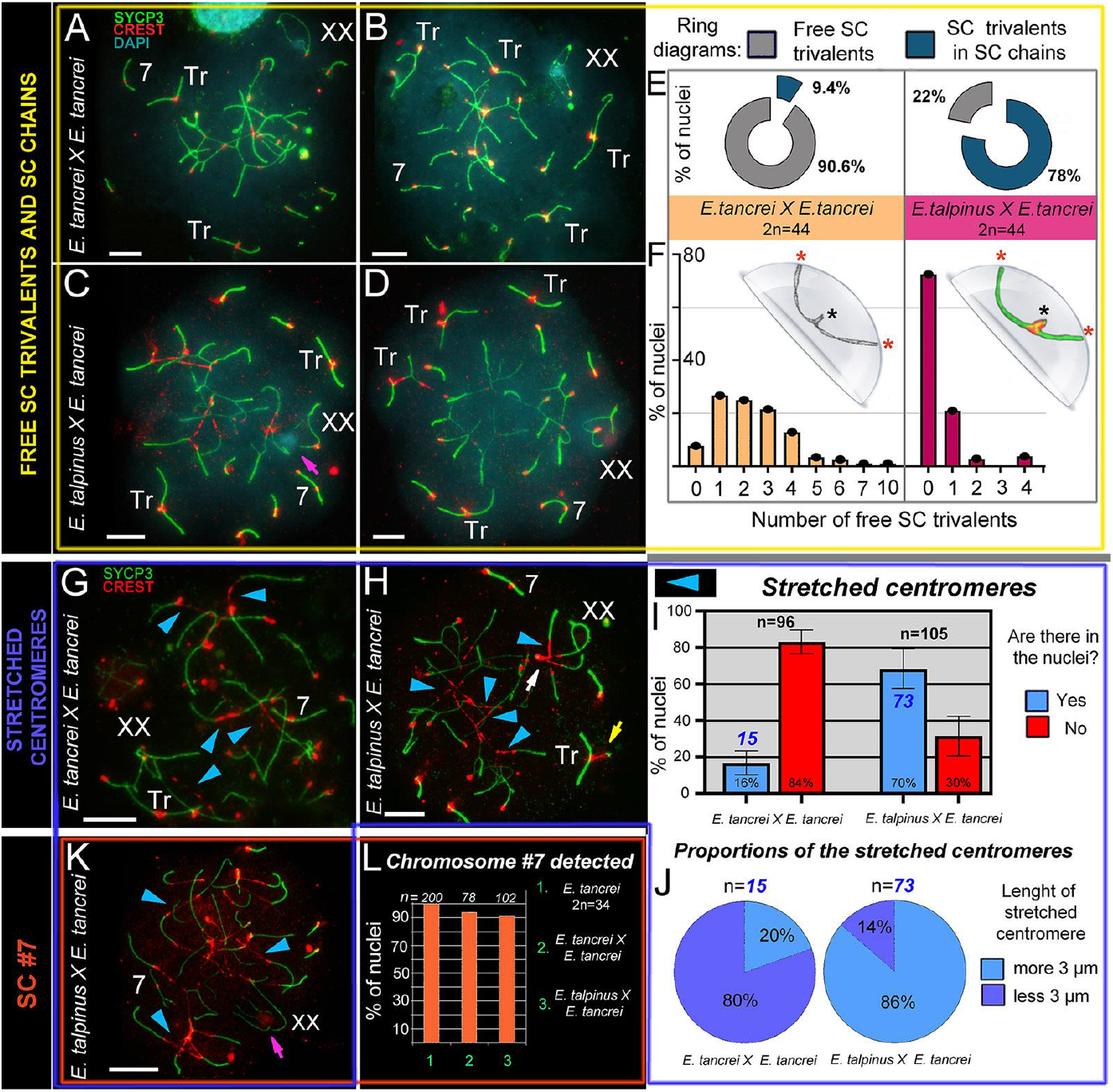
SC trivalents and stretched centromeres in *Ellobius* hybrids. **(A–F)** SC trivalents, free and in chains, in 44-chromosome intraspecific (A, B) and interspecific (C, D) hybrids. Both hybrids had variations in the number of free closed trivalents (see A–D). Fewer free closed SC trivalents in the nuclei indicated that more open SC trivalents were in the chains. The number of free trivalents in hybrid nuclei was significantly different: 90.6% in intraspecific (n = 99) and 22% in interspecific (n = 102) hybrids (see ring diagrams in E and see Table S1). There were typically 1–4 free SC trivalents identified in most nuclei of intraspecific hybrids (see the light orange columns of the bar chart in F). As a rule, there were no free SC trivalents in most nuclei of interspecific hybrids (see the dark-red columns of the bar chart in F). Free closed SC trivalents simulated in the hemisphere (F): Black stars show proximal telomeres of acrocentrics, and red stars show distal telomeres of metacentrics and acrocentrics of SC trivalents. **(G–K)** Stretched centromeres in SC trivalents (blue arrowheads in G, H, K). The prevalence of the stretched centromeres in hybrid nuclei was variable: 16% (15 nuclei from n = 96) in intraspecific and 70% (73 from n = 105) in interspecific hybrid (see bar chart in I; Table S1 indicates significant differences). The length of the stretched centromeres was different; there were two notable groups: (1) < 3 μm and (2) > 3 μm. Group (1) prevailed in the intraspecific hybrid, while group (2) was more prevalent in the interspecific hybrid (see ring diagrams in J). **(L)** Prevalence of chromosome #7 detected in the parent (2n = 34) and hybrids. Chromosome #7 was clearly identified in all nuclei (see A–D, G, H, and K). Parents (2n = 34) always had chromosome #7 (without chromosome associations). Both types of hybrids had chromosome #7 in the vast majority of pachytene nuclei (see the columns of the bar chart in L). Pink arrows show associations of the sex (XX) bivalents (C, K). The white arrow shows atypical lengthening of one of the XX bivalent synaptic sites (H). The yellow arrow shows atypical prolonging of short arms of acrocentrics of the SC trivalent (H). The nucleus in K is presented in Fig S6A–G). Scale bars represent 5 μm.

The interspecific hybrid also exhibited atypically large XX bivalents (Fig 2G) and ring univalents at the pachytene stage (Fig S4I). The intraspecific hybrid showed a triple synapsis in the region of the short arms of the SC trivalent (Fig S3G).

Associations of the XX bivalent with autosomes were rare and only observed in two forms of *E. tancrei* nuclei: 5.9% (2n = 54) and 2.25% (2n = 34). In both F1 hybrids, associations of the sex bivalent with autosomes/trivalents were much more common: 50.37% (intraspecific hybrid) and 45.34% (interspecific hybrid) of the nuclei had these features (Fig 2K, Table S1[4]). Examples of XX bivalent association with SC trivalents are presented in several of the figures (intraspecific: Fig 2D, 2E and 2F, S3A, S3B, S3C, and S3E; interspecific: Fig 2H, S4A, and S4B). It should be emphasized that in interspecific hybrids, there were associations of one of the X axes with an autosome or trivalent by a thin CREST-positive linear link (Fig S4B and S4D), as well as an association through a ChB (Fig 3C).

### Free closed SC trivalents and open SC trivalents in chains

The intra- and interspecific hybrids differed in the number of formed free closed SC trivalents. Based on the karyotypes, we assumed that at the pachytene stage of both hybrids, 10 metacentrics and 20 acrocentrics should have formed 10 SC trivalents. However, in intraspecific hybrids, 1–4 free closed SC trivalents were most often found (Fig 3A, 3B, 3G, and S3F). It should be emphasized that in single nuclei, we found 7 and 10 SC trivalents (Fig 3E, 3F, S3C and S3H). In one of the animals, there were closed SC trivalents with SYCP3- and AgNO3-positive dense material in the region of the short arms of the acrocentrics (Fig S3H, S3H’, S3H”, S3I, S3J, S3J’, and S3K) or in the pericentromeric region of SC trivalents (Fig S3L and S3L’). Open SC trivalents formed SC chains due to heterosynapsis between the short arms of acrocentrics (Fig S3A and S3B).

Free closed SC trivalents, as a rule, were not detected in spermatocytes of interspecific hybrids (Fig 3E, 3F, 3K, S4A, S4J, S4K, and S4L, Table S1[5]), i.e., they remained open and embedded in SC chains. There was 1 closed SC trivalent in 21% of nuclei (Fig 3H), as well as in squashes (Fig S8). There were 2 (Fig 3C) or 4 (Fig 3D) SC trivalents in single cells (Fig 3F). In the chains of the SC trivalents, the centromeric sites of the metacentrics were strongly stretched (see next section).

### Stretched centromeres in SC trivalents and their participation in SC trivalents chains

In the nuclei of cells from both hybrids, there was stretching of the centromeric regions of the metacentrics of the SC trivalents (intraspecific: Fig 3A and 3G; interspecific: Fig 3C, 3D, 3H, and 3K). However, we only observed this phenomenon in 16% of pachytene cells from intraspecific hybrids, and their length was less than 3 μm (Fig 3I and 3J). By contrast, in interspecific hybrids, we observed centromere stretching in 70% of spermatocytes. In most cases the length was > 3 μm (Fig 3I and 3J, Table S1[6]), and in some nuclei it reached ≥ 10 μm (for example, Fig 3K). In interspecific hybrids, the centromeric region of the metacentrics was strongly stretched (Fig 3H, 3K, S4B, and S4F). There was trivalent centromere stretching in zygotene spreads (Fig S5) and pachytene squashes (Fig S8). Squashes with preserved three-dimensional nuclear space confirmed that stretched centromeres were not an artifact of the spreading technique (Fig S8).

Of note, we detected centromere stretching in the acrocentrics of the SC trivalents, although less frequently than in metacentrics (Fig S6). This feature was shown in SC trivalents, which were part of SC chains (Fig. S6A, S6B, S6C, S6D, S6E, S6F, S6G, and S6I).

Thus, a comparison of the chains of SC trivalents allowed us to establish some differences between hybrids. In intraspecific hybrids, the connection between trivalents occurred due to the short arms of acrocentrics (heterosynapsis). In interspecific hybrids, the formation of such SC chains was accompanied by a strong stretching of the centromeric regions of the metacentrics and acrocentrics in SC trivalents, and neighboring SC trivalents can join due to the stretched centromeric regions of the acrocentrics (shown schematically in Fig S7 for both hybrids).

In general, three main types of SC trivalents can be distinguished in mole vole hybrids (Fig 4). It should be noted that we identified centromere stretching by immunostaining with both polyclonal and monoclonal antibodies to centromere proteins obtained from different manufacturers (Fig S4A, S4J, S4K and S4L).

**Fig 4.**
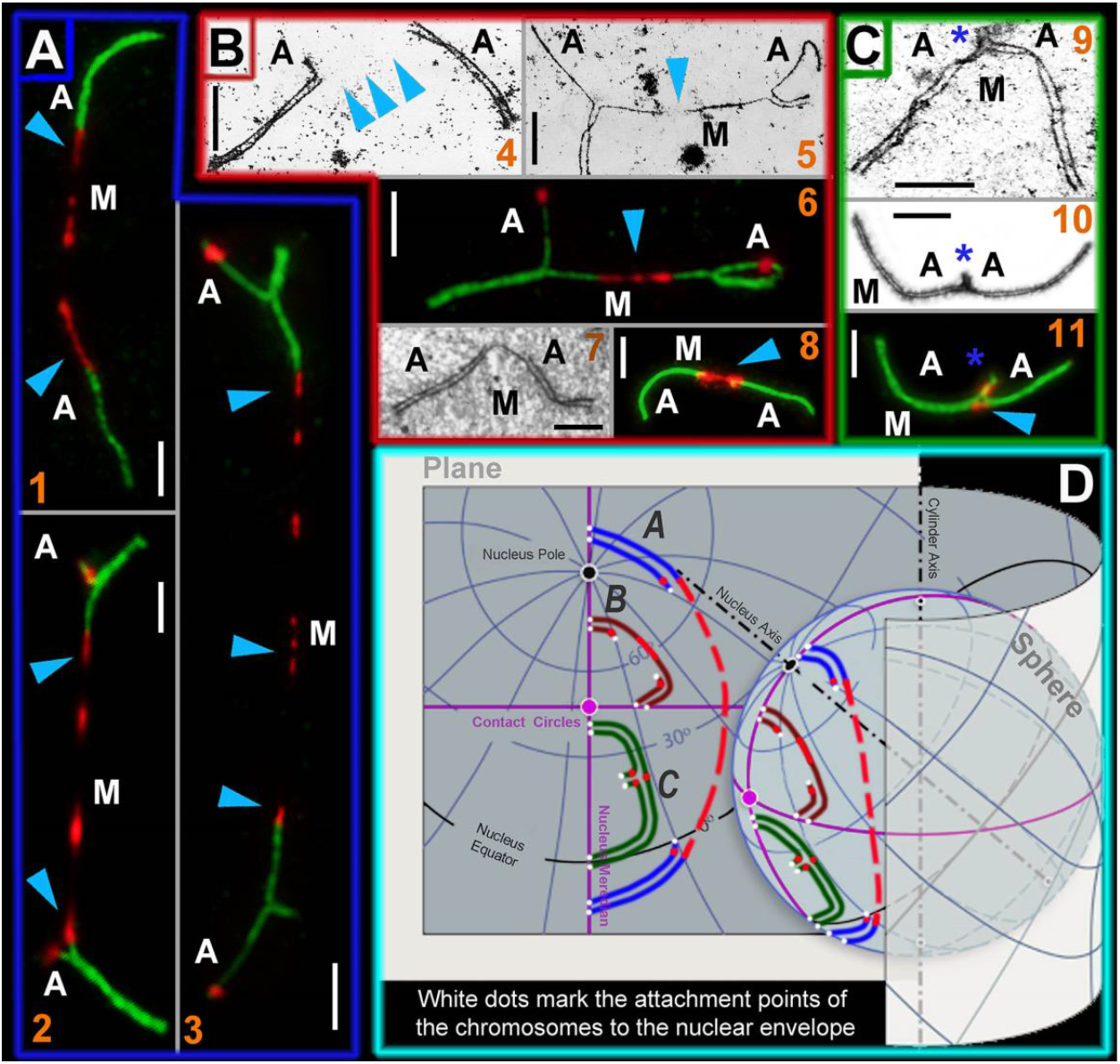
SC trivalent types of the *Ellobius* hybrids and their simulated location on a plane and a sphere. Light micrographs after immunostaining (1, 2, 3, 6, 8, and 11): SCs were immunostained with antibodies against SYCP3 (green) and centromeres—with an antibody to kinetochores (CREST, red). Electron micrographs after AgNO3-staining (4, 5, 7, 9, and 10): Blue arrowheads show centromeric regions of metacentrics in SC trivalents. M indicates metacentric and A indicates acrocentric. **(A)** SC trivalents elongated by centromeric region stretching in spermatocytes of the *E talpinus* × *E tancrei* interspecific hybrid. The length of the centromeric region of metacentrics in such SC trivalents is > 3 μm. (1) SC trivalents in which short arms were not visible: If there were no immunostained centromeric regions, then the SC trivalents would be identified as two pseudobivalents. (2) SC trivalents with short arms of acrocentrics. (3) SC trivalents with very stretched centromere. As a rule, such SC trivalents stretch across the length of the entire nucleus (from one side to the other). **(B)** SC trivalents with short stretched centromeric regions (< 3 μm in length). This type of SC trivalent was found in all hybrids. (4) Two pseudobivalents that are part of the SC trivalent. (5) The metacentric in the SC trivalent has a gap, in which there is probably a stretched centromeric region. (6) SC trivalents like in 5; the stretched centromere of the metacentric are visible. (7 and 8) Short SC trivalents with faintly distinguished short arms of acrocentrics and not very stretched centromeres. **(C)** Closed SC trivalents with clearly visible short arms of acrocentrics (blue stars) and dot-like centromeres (9–11): This type of SC trivalent is found in all hybrids. **(D)** Simulation of SC trivalents on a plane and a sphere. SC trivalents of *A* type (see A) have far-reaching attachment points to the nuclear envelope and long stretched centromeric region with gaps. SC trivalents of *B* and *C* types (see panels B and C) are more compact and do not have long stretched centromeres. Scale bars represent 5 μm.

### Bivalent #7 behavior

Chromosome #7 is an excellent karyotypic marker, and it can be used to identify both intra- and interspecific hybrids. Based on previous research, the submetacentric in *E. tancrei* is formed due to the neocentromere formation established by SCs analyses (Matveevsky, 2011a, 2011b) and then supplemented by Zoo-FISH data (Bakloushinskaya et al., 2012). In interspecific hybrids, this chromosome pair appears as an SC bivalent with two centromeres, because an acrocentric homolog (*E. talpinus*) and a submetacentric homolog (*E. tancrei*) entered into synapsis. In the vast majority of nuclei in both hybrids, SC #7 was easily identified (Fig 2 and 3, Table S1[7]). There was only a clear association of the SC chromosome #7 bivalent, or rather its acrocentric homolog, with the SC trivalent (Fig S4E).

### Chromosome recombination

We studied meiotic recombination in *Ellobius* parental species and hybrids using the MLH1 protein, a marker of crossovers. The average number of MLH1 foci per nucleus is an important recombination parameter. It can be used to compare closely related species, in homozygous and heterozygous forms (Bhattacharyya et al., 2013), as well as to examine recombination in larger taxa (Segura et al., 2013). In the latter case, the haploid number of autosomes (NFha) and the haploid number of autosomal arms (NFha.a) are considered.

The number of MLH1 foci per nucleus (M ± SD) was 23.48 ± 3.3 in *E. talpinus* (2n = 54, NF = 54; NFha = 26, NHha.a = 26), 23.1 ± 3.1 in *E. tancrei* (2n = 54, NF = 56; NFha = 26, NHha.a = 27), and 22.8 ± 3.7 in *E. tancrei* (2n = 34, NF = 56; NFha = 16, NHha.a = 27) (Fig 5A–C). These rates were not significantly different among species (Table S1). The number of MLH1 foci in all species was slightly lower than the NFha.a values. Thus, not every chromosome arm has at least one MLH1 signal. We previously described detection of MLH1 signals in the XX bivalent; the rate was 46% for *E. talpinus* and 65% for *E. tancrei* (2n = 54) (Matveevsky et al., 2016).

**Fig 5.**
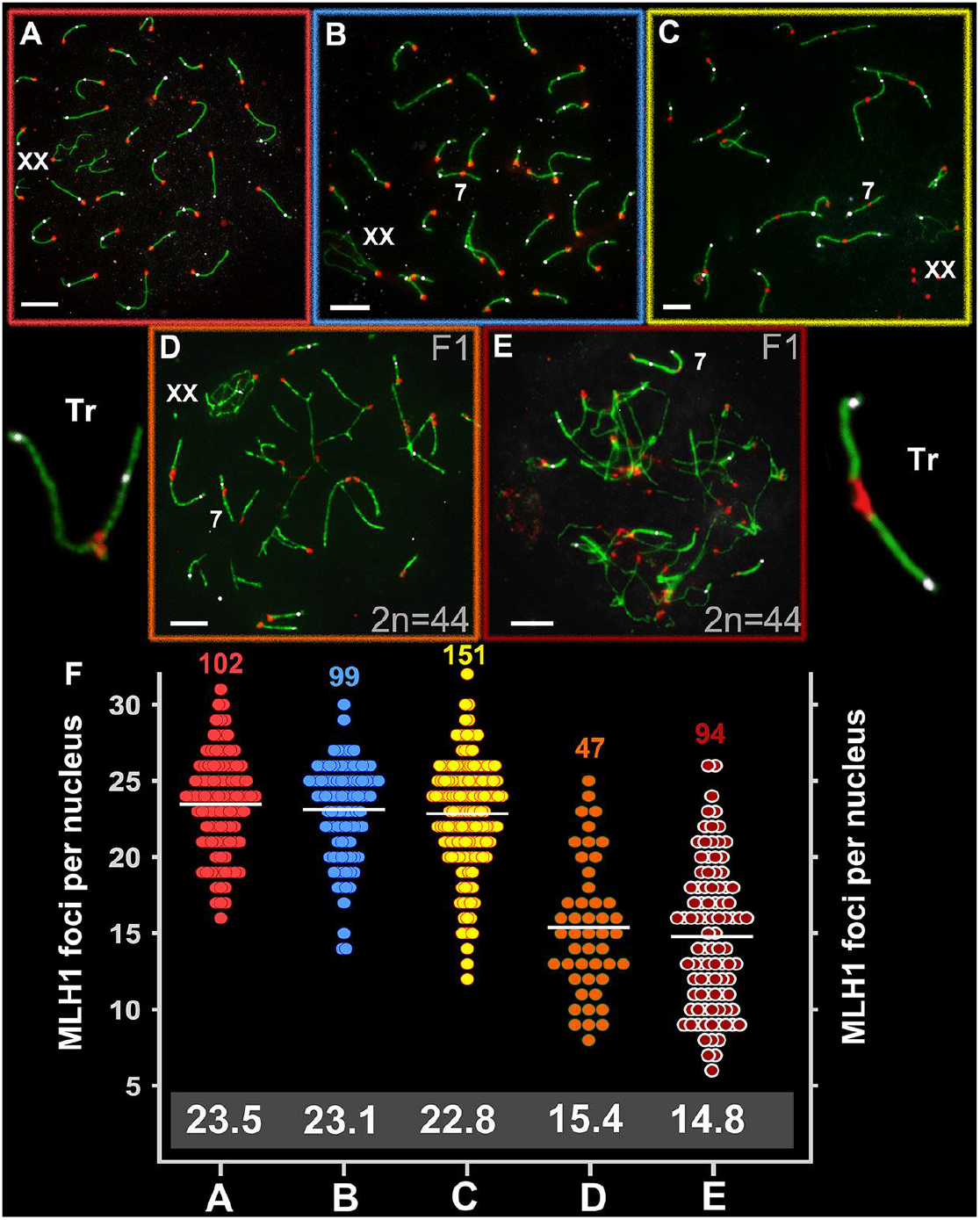
Recombination in *Ellobius* species and hybrid males. Axial elements were identified using anti-SYCP3 antibodies (green), kinetochores using CREST antibodies (red), and anti-MLH1 (white) was used as a marker of recombination nodules. **(A)** *E talpinus* (2n = 54, NF = 54); **(B)** *E. tancrei* (2n = 54, NF = 56); **(C)** *E. tancrei* (2n = 34, NF = 54); (D) F1 *E. tancrei* × *E. tancrei* hybrid; **(E)** F1 hybrid *E. talpinus* × *E. tancrei* hybrid. **(F)** Comparison of MLH1 foci per nucleus in mole voles: The numbers above the dot diagrams correspond to the number of counted cells. White numbers in a gray frame correspond to the average number of MLH1 foci. Table S1 presents details on significant statistical differences between species and hybrids. Scale bars represent 5 μm.

The average number of MLH1 signals was 15.4 ± 4.4 for the intraspecific hybrid (2n = 44, NF = 56; NFha = 21, NHha.a = 27) and 14.8 ± 4.7 for the interspecific hybrid (2n = 44, NF = 55; NFha = 21, NHha.a = 26.5) (Fig 5D and 5E). These rates differed significantly between the hybrids and compared with the parents (Table S1) and were significantly lower than the NHha.a number. A decrease in the recombination level in hybrid spermatocytes was due to the formation of chains of SC trivalents, where the arms of acrocentrics did not have a complete synapse with the arms of metacentrics.

## Discussion

Rb translocations affect the genome by modifying gene positions and altering recombination during meiosis. Overall, our findings present experimental evidence supporting assumptions that chromosomal rearrangements redesign a genome and may contribute to speciation due to meiotic alterations in hybrids. We have demonstrated that hybrids with the same diploid number and identical chromosome combination could be sterile (interspecific) or have reduced fertility (intraspecific), a condition that is in line with the behavior of chromosomes in meiosis. Despite the fact that both types of hybrids had similar irregularities during prophase I, there were different synaptic patterns in SC trivalents, especially specifics of centromeric segments. We hypothesize that these differences originated due to altered meiotic architecture and could to be responsible for the species divergence.

### Chromosome synapsis instability and reduced recombination in hybrids

There is a huge amount of data and knowledge that allows us to unambiguously state that hybrids possess distinct patterns of chromosomal synapsis and recombination compared with their parents, which have a significant evolutionary output (King, 1993, Arnold, 1997, Abbott et al., 2016). Intra- and interspecific mole vole hybrids with 10 trivalents showed variation in fertility, as has been shown for heterozygous lemurs. Indeed, the first known case of hybrids with numerous Rb chromosomes was described for lemurs (Moses et al., 1979). At that time, researchers believed that hybridization occurred between lemur subspecies (Ratomponirina et al., 1988). According to the modern taxonomy (Wilson and Reeader, 2005, Schwitzer et al., 2013), interspecific lemur hybrids were studied in 1988. Lemurs with 3–6 trivalents had fertility similar to their parents, while lemurs with 8 trivalents had reduced fertility (Moses et al., 1979; Ratomponirina et al., 1988).

The first group of lemurs did not have any associations of bivalents and trivalents with each other and with XY. The second group regularly demonstrated chains of SC trivalents, combined by heterosynapsis of the short arms of acrocentrics, and association with the sex bivalent (Ratomponirina et al., 1988). A large number of trivalents were likely unable to complete synapsis in time, and this deficit could lead to aberrant chromosome segregation, arrest of cells at M1 or M2 stages, germ cell aneuploidy, and decreased fertility. We observed the synapsis delay for SC trivalents in intraspecific mole vole hybrids; this finding is consistent with previous studies (Kolomiets et al., 1985, Bogdanov et al., 1986). However, in contrast to lemur hybrids, in intraspecific mole vole hybrids, there were often gaps in the axial elements of the metacentrics in open SC trivalents (Kolomiets et al., 1985; Bogdanov et al., 1986). Based on immunostaining, now we interpret such specific gaps as stretching of the centromeric regions.

*Mus musculus domesticus* that were heterozygous for Rb metacentrics were also highly variable in their fertility (de Boer and de Jong, 1989; Hauffe and Searle, 1998). Thus, in some intraspecific mice hybrids, for example, heterozygous for three Rb translocations, there was no decrease of fertility, although there were “XY–trivalent” and “trivalent-trivalent” associations (Grao et al., 1989). In other cases, comparative analysis revealed differences in fertility level due to distinct karyotype structures: a slight decrease (4 SC trivalents) and a significant decline in fertility (7 SC trivalents) (Wallace et al., 2002). Possibly, the difference was due to higher associations of SC trivalents with sex XY bivalent and associations of trivalents with each other in animals of the second group.

In other experiments, fertile hybrids heterozygous for 8 Rb translocations had open and closed SC trivalents (Manterola et al., 2009). The authors suggested that the meiotic progression of cells with an asynaptic area of SC trivalents was due to the insignificant genetic value of the unsynapsed chromatin regions, the inactivation of which did not lead to the activation of the pachytene arrest program.

Hybridization of different forms of the musk shrew (*Suncus murinus*) could lead to the formation of 5 SC trivalents in meiocytes. Some of the F1 hybrids were sterile, while others were fertile, depending on different parental combinations. Researchers have suggested that genetic factors play a crucial role in determining fertility/sterility (Borodin et al., 1998; Rogatcheva et al., 2000). In the case of mole vole hybrids, animals regularly had impaired fertility (intraspecific) or were sterile (interspecific), although this fact does not exclude the effect of unknown genetic factors in determining the fertility level.

Alterations in recombination might be crucial for species evolution (Rieseberg, 2001, Livingstone and Rieseberg, 2004). Physical problems in the synapsis of rearranged homologs can restrict the formation of recombination sites (Cattanach and Moseley, 1973, Davisson and Akeson, 1993), which can lead to univalence, unbalanced chromosome segregation and selection of germ cells (Baker et al., 1996, Eaker et al., 2001, 2002). Comparison of distinct mole vole hybrids reliably demonstrated that recombination was reduced in hybrids, a phenomenon that was most likely correlated with a delay in synapsis of SC trivalents or their physical stretching. It is likely that some of the achiasmatic chromosomes in trivalents can be incorrectly segregated, and such cells will be eliminated in meiotic checkpoints (Morelli and Cohen, 2005).

### Centromere identity and stretched centromeres of the SC trivalents

Centromeres are unique chromosomal regions that organize the assembly of the kinetochore, a large multiprotein complex that allows chromosomes to attach to spindle microtubules and move during mitotic and meiotic cell division (Cheeseman, 2014, Talbert and Henikoff, 2020). There is no universal DNA sequence responsible for centromere formation. This fact, and the emergence of new centromeres, led to the hypothesis that centromeres are determined by accumulation of tandem repeats (satellite DNA) (Henikoff et al., 2001, Garagna et al., 2002, Plohl et al., 2008) and retrotransposons (Ferreri et al., 2011, Chang et al., 2019) alongside with epigenetic modifications such as histone variants (Amor and Choo, 2002, Dalal, 2009). The centromere-specific variant of histone H3, CENPA, is a platform for protein assembly at the kinetochore (Bernard et al., 2009, Musacchio and Desai, 2017). The rapid diversification of centromeres has been suspected to lead to reproductive isolation between species (Henikoff et al., 2001). The position of the centromere is easily identified by routine staining; in immunostaining centromeres typically appear as one-spotted signals within chromosomes.

In mole voles, closed SC trivalents developed three contact points with the nuclear envelope: (1) the attachment point of the proximal ends of acrocentrics, formed by their short arms, and (2 and 3) two attachment points of the distal telomeres of metacentrics and acrocentrics (see Fig 3F). As prophase I progresses, the attachment points move away from each other; therefore, SC trivalents can have different spatial configurations (Berrios et al., 2014). However, open SC trivalents must be associated with the nuclear envelope at four points. In addition, acrocentrics were ectopically connected by short SCs with neighboring acrocentrics. There were multiple interlockings in interspecific mole vole hybrids. All these specifics caused the strongest tension of chromosomes involved in SC trivalents, stretching of the centromeric regions of the Rb metacentrics, and in some cases, the centromeric regions of the acrocentrics.

We used different antibodies against kinetochore proteins to identify centromeric regions of meiotic chromosomes in mole vole parental species and hybrids (Fig S4). As noted above, we observed 3 types of SC trivalents in hybrid spermatocytes (Fig 4). Thus, the distance between the attachment points determined the ultrastructural organization of SC trivalents. These attachment points were most clearly manifested in closed free SC trivalents (Fig 4C). If the distance between the attachment points of the SC trivalent to the nuclear envelope was markedly greater than the metacentric length, then the metacentric axis undergoes strong stretching, and this did not allow the formation of a continuous axial element. Therefore, electron microscopic examination revealed a gap in the structure of the stretched axial element of the metacentric (Fig 4B). However, when immunostaining with antibodies to kinetochore proteins (ACA, CREST, and CENPA), there were no gaps, and a centromeric linear structure was visible in this area (Fig 4B). If the attachment points were located very far from each other (on different sides of the nucleus), then the centromeric region was hyperstretched up to gaps (Fig 4A).

Such stretched centromeres are intriguing because no similar regions have been found within SC trivalents before the present study. It is possible that these centromeric stretches are associated with the structural specifics of the pericentromeric chromatin. Chromatin in the centromeric region has the classic epigenetic marks of constitutive heterochromatin: H3K9me2, H3K9me3, H3K27me3, and H4K20me3 (Richards and Elgin, 2002; Nishibuchi and Dejardin, 2017). For example, in the study of unfolded pericentromeric heterochromatin (prekinetochores) in interphase, chromosomes subjected to stretching by TEEN buffer, there was alternation of the CENPA and H3K9me3 subdomains with the gaps (Ribeiro et al., 2010). If any natural or artificial extension does not lead to structural breaks, it likely entails a linear spatial unfolding of the structural components of pericentromeric heterochromatin. This phenomenon probably explains why we saw an alternating change of centromeric points, centromeric lines, and gaps in the metacentrics of SC trivalents. This feature might indicate high plasticity of mole vole pericentromeric heterochromatin.

Stretching of the centromeric regions of acrocentrics between SC trivalents is even more mysterious. This phenomenon was rarer than centromeric stretching of metacentrics. We suppose that this may also be due to a special stretching of the pericentromeric heterochromatin of the acrocentric short arms associated in chains of several SC trivalents. Moreover, heterochromatin was involved in the nonhomologous synapsis of the short arms of the acrocentrics of SC trivalents and in binding to the nuclear envelope in intraspecific hybrids (Fig S9B and S9C). The centromeric associations of SC trivalents are of particular interest (see “Nuclear architecture: simulated chromosome configurations in mole vole pachytene spermatocytes “ below).

In general, the presence of stretched centromeric regions may indicate specific properties of centromeres in the genus *Ellobius*. Stretched centromeres in bivalents as a likely result of Rb chromosome fusion have been identified in African pygmy mouse (Gil-Fernández et al., 2020). Centromeric satellites in animals and plants undergo rapid evolution (Henikoff et al., 2001), and they may differ even in closely related species (Meštrović et al., 1998, Lee et al., 2005, Talbert et al., 2018). These differences have been explained by the “library hypothesis” (Salser et al. 1976): An ancestral form has an initial satellite pool (“library” of satellites), which in different evolutionary lineages manifests in various patterns (in quantity and quality), thus forming species-specific satellite profiles (Miklos et al., 1985). The hypothesis is supported by some examples (see Smalec et al., 2019). In addition, it has been established that closely related species may differ in their retrotransposons. A classic example is a well-studied kangaroo. In interspecific kangaroo hybrids, there was centromere destabilization, which was caused by the activation of resident retroelements (in this case, kangaroo endogenous retrovirus [KERV]) (O’Neill et al., 1998, Metcalfe et al., 2007, Ferreri et al., 2011).

It would seem that such important chromosome elements like centromeres should be conserved, but they exhibit incredible structural and functional variability and dynamic evolution and are hotspots for chromosomal rearrangements (Brown and O’Neill, 2010, Brown et al., 2012). The main remaining issues include the question of the true reasons for the centromeric region stretching in *Ellobius* hybrids as well as the molecular mechanisms of their striking plasticity. The study of centromeric satellites and retrotransposons in the *Ellobius* genus will be promising. It is obvious that additional detailed study of centromeric regions in *Ellobius*, including stretched centromeres and different centromeric localization of the heteromorphic chromosome #7 pair in interspecific hybrids, will be necessary. The centromere features established here and data from previous works (Matveevsky, 2011a, 2011b, Bakloushinskaya et al., 2012, Matveevsky et al., 2017) may suggest that the centromeres in mole voles, if not a driver of chromosomal evolution, are essential for karyotypic divergence.

### Chromosome #7: implications from the hybrid meiotic nuclei studies

As mentioned above, the parental 54-chromosome karyotypes of *E. talpinus* and *E. tancrei* differ in only one pair of chromosomes (#7). In these species, chromosome #7 pairs were identical in G-bands, but in *E. talpinus* it is an acrocentric, and in *E. tancrei* it is a submetacentric. For a long time, it was assumed that the submetacentric emergence was due to pericentric inversion (Lyapunova and Vorontsov, 1978). However, the SC study in interspecific hybrids clearly showed that this chromosome pair is completely synapsed without inversion loops, forming a full-length SC along the entire chromosome at the early-to-mid pachytene stage. We previously suggested that the submetacentric in *E. tancrei* emerged through the neocentromere formation (Matveevsky, 2011a, 2011b). This assumption was confirmed by additional Zoo-FISH data (Bakloushinskaya et al., 2012), and later by immuno-identification of the central element of SCs and recombination nodules (Matveevsky et al., 2017).

The chromosome #7 pair was used as a marker bivalent in the SC analysis. In interspecific hybrids, this was the only heteromorphic SC bivalent with two centromeres located at a distance from each other. Of note, we discovered another remarkable property of the SC #7. In both hybrids, bivalent #7 practically does not participate in chromosome associations: There was only one association of this chromosome with the open SC trivalent (Fig S4E). We assume that the inertness of chromosome #7 in meiosis may be due to the absence or extremely low content of constitutive heterochromatin. This was confirmed using antibodies to histone H3K9me3, a marker of constitutive heterochromatin, both in bivalent #7 in *E. tancrei* (unpublished) and in a sibling species *E. alaicus* (Matveevsky et al., 2020).

Thus, the ancestors of modern *E. tancrei* and *E. alaicus*, concomitant with the emergence of the neocentromeric submetacentric chromosome #7 pair, probably acquired chromosomal instability with the formation of various karyotypic forms, which may indicate some causal link between these events. Perhaps one of them could become an evolutionary trigger, which entailed a chain of genetic and/or cytogenetic changes in the mole vole karyotype.

### Nuclear architecture: simulated chromosome configurations in mole vole pachytene spermatocytes

The organization of the internal contents is not random in interphase (Cremer et al., 1993, Parada and Misteli, 2002) and meiotic (Foster et al., 2005) nuclei; therefore, it is customary to speak of nuclear architecture (Cremer and Cremer, 2010) or intranuclear landscape (Cremer et al., 2006). The “chromosome territory” concept is considered to be generally accepted (Cremer and Cremer, 2001). Chromosome territories are spatial domains of different sizes that specifically occupy a certain volume in the nucleus (Misteli, 2008). The position of chromosomes in the nucleus can be preserved in closely related taxa (Tanabe et al., 2002), a phenomenon called “phylogenetic memory” (Mora et al., 2006). Heterochromatic compartments play a significant role in the nucleus content (Politz et al., 2016). However, gene mutations and chromatin and chromosomal rearrangements can change the intranuclear organization, which can lead to diseases and the appearance of various abnormal manifestations (Garagna et al., 2001, Oberdoerffer and Sinclair, 2007, Lammerding, 2011, Fournier et al., 2010, Berrios et al., 2018, Abdelhedi et al., 2019).

There is also a point of view, according to which the correct formation of chromosome territories diminishes the translocation potential of the cells (Rosin et al., 2019). The organization of interphase and meiotic nuclei has significant differences (Alsheimer et al., 1999). An SC forms a specific chromatin pattern (Hernandez-Hernandez et al., 2009) and interacts with the nuclear envelope through a special Sun-KASH system (Alsheimer, 2009). The presence of asynaptic chromosome regions in prophase I initiates a meiotic silencing of unsynapsed chromatin (MSUC) (Schimenti, 2005; Turner et al., 2005). Nuclear architecture of prophase I meiocytes is specific for each species (Berrios et al., 1999, 2004, Berrios, 2017).

As a result, the nuclear architecture in meiotic prophase I is determined by the SC structure and dynamics, types of chromosomes (single-armed or bi-armed), their length, the heterochromatin amount, the specificity of centromeric regions, the “chromosome–nuclear envelop” interactions, the ability to form chromocenters and nucleoli, and sex chromosome organization and behavior (Berrios, 2017). It should be emphasized that if the parental genomes differ significantly, then complex chromosome compounds are formed in hybrid and mutant meiotic nuclei (for example, Johannisson and Winking, 1994, Ribagorda et al., 2019, Matveevsky and Kolomiets, 2011, 2016), and the processes of repair, recombination, and meiotic silencing are disrupted (for example, Bhattacharyya et al., 2013, Turner, 2007, Burgoyne et al., 2009, Homolka et al., 2007), which can cause an imbalance in the nuclear architecture.

In two hybrid mole vole groups, a different number of free closed SC trivalents formed. Pericentromeric heterochromatin likely played an important role in the formation of closed trivalent configurations, as described in heterozygous mice (Berrios et al., 2014). Thus, on the three-dimensional nuclei of interspecific mole vole hybrids, there were closed SC trivalents, which were attached by the short arms of acrocentrics to the nuclear envelope, and open SC trivalents with stretched centromeric regions (CREST cloud around) (Fig S8). CREST antibodies can non-specifically immunostain heterochromatic regions and heterochromatin-like structures, for example, ChBs (Fig 2H and 3D). The same specificity was revealed during the electron microscopic examination of the intraspecific heterozygous spermatocytes. At the attachment site of the short arms of the SC trivalent, a cloud of electron-dense material formed, associated with the nuclear envelope, which was usually interpreted as heterochromatin mass (Fig S9, schemes of hemispheres in 3F).

SC trivalent chains were demonstrated in intraspecific mole vole hybrids (Kolomiets et al., 1985; Bogdanov et al., 1986). Such ectopic associations formed due to the heterochromatic contacts of the short arms of acrocentrics of two different open SC trivalents (Fig S7), as had been noted earlier (Kolomiets et al., 1986). Chromosome associations can be determined by H3K9me3 immunodetection (Berrios et al., 2014). This protein is believed to mark DAPI-positive chromocenters (for example, Berrios et al., 2010). However, chromocenters in all mole voles were usually not detected by DAPI staining. Nevertheless, we found that acrocentric chromosomes were grouped by their pericentromeric regions around the H3K9me3 domains in the sibling species *E. alaicus* (Matveevsky et al., 2020). We saw a similar grouped position of acrocentrics in *E. talpinus* (Fig 2A). In *E. tancrei* (2n = 34), the centromeric sites of the acrocentrics were also located around the H3K9me3 clouds, while these sites of the Rb metacentrics had a more linear form of the H3K9me3 signals, as seen in the micrographs (Fig S9A, S9A’).

Combining the results on the ultrastructure and behavior of SCs and SC trivalents in spreads and squashes, we present these data as three-dimensional simulations of chromosome configurations in the meiotic nuclei of mole vole parental species and hybrids (Fig 6).

**Fig 6.**
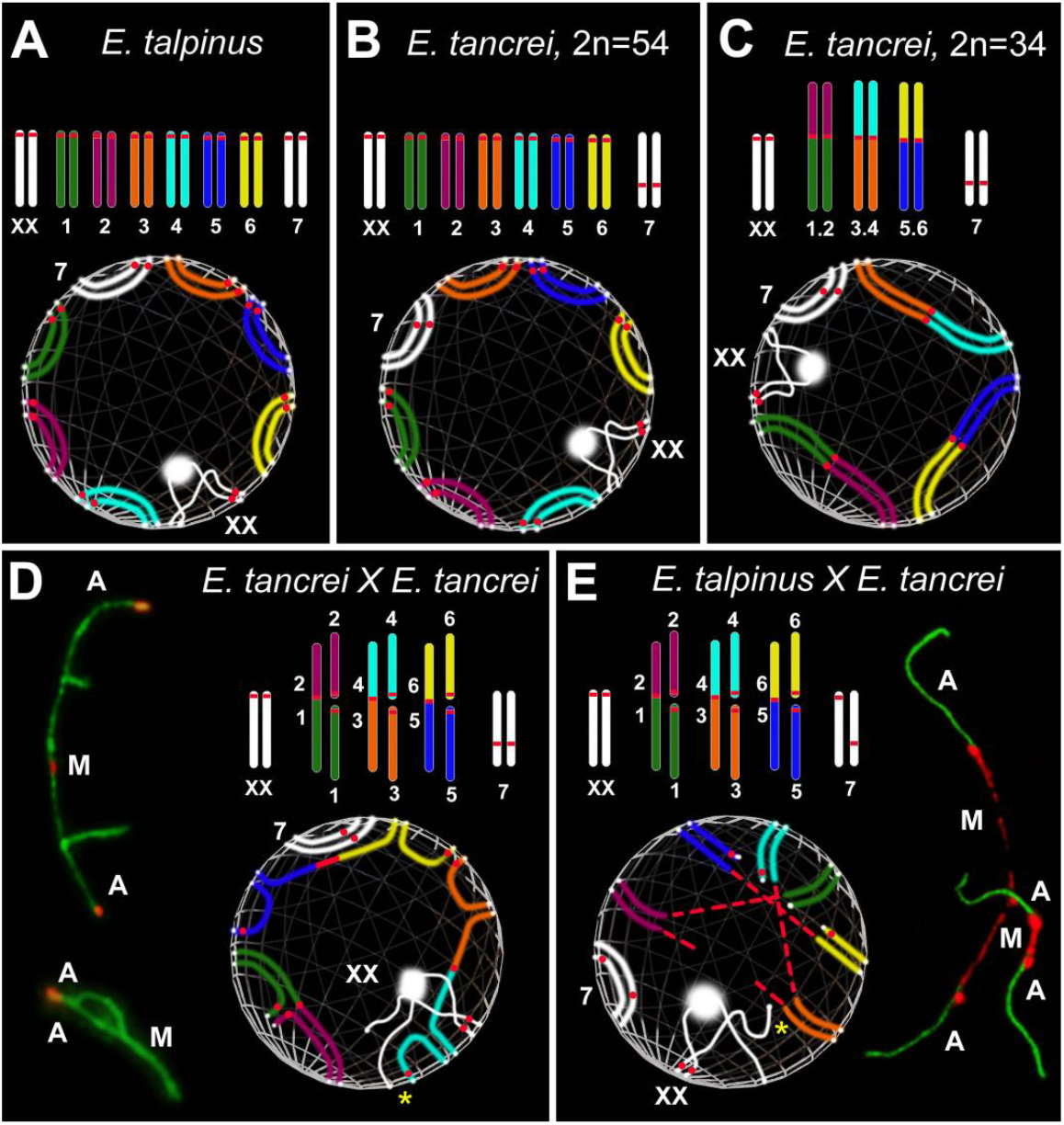
Simulation of chromosome synapsis and interactions of SC trivalents in pachytene nuclei of *Ellobius* species and hybrids. A partial set of chromosomes and lower—inside the spheres—features of the configuration of the axial elements of these chromosomes during synapsis at the pachytene stage are presented in each part of the figure. White dots at chromosome ends mark the attachment points of the chromosomes to the nuclear envelope. **(A**, **B)** *Chromosomes:* Seven pairs of autosomes and a pair of isomorphic sex (XX) chromosomes in A and B. The chromosome sets differ from each other only in the chromosome 7 pair: acrocentric in *Ellobius talpinus* (2n = 54), neocentromeric submetacentric in *Ellobius tancrei* (2n = 54). *Spheres:* The positions of SCs over the nuclear envelope in the pachytene nuclei in A and B are different, because it is assumed that chromosome territories can be distinguished between two species (Berrios et al., 2014, 2017). **(C)** *Chromosomes:* Conditional karyotype of *E. tancrei* (2n = 34) consists of three pairs of homologous Robertsonian (Rb) metacentrics (1.2, 3.4 and 5.6), a submetacentric chromosome #7 pair and sex (XX) chromosomes. *Sphere:* The chromosomal arms of metacentrics do not always occupy the ancestral position (compare spheres C, B, and A), because Rb metacentric formation causes a shift in the chromosome territories in the meiotic nucleus (see Garagna et al., 2001, Berrios et al., 2014). **(D)** *Chromosomes:* Karyotype of the intraspecific hybrid *E. tancrei* obtained from crossing two forms (2n = 54 and 2n = 34). The scheme of partial karyotype shows three Rb metacentrics (1.2, 3.4, 5.6) and six acrocentrics (1–6). *Sphere:* Three Rb metacentrics and acrocentrics form three SC trivalents. They are attached to the nuclear envelope and part of them remains open for a long time at the mid pachytene. The synapsis of chromosomes in all SC trivalents is gradually completed to the diplotene, which will ensure their correct segregation in the future. Open and closed SC trivalents are presented on the left. **(E)** *Chromosomes:* Karyotype of the sterile interspecific hybrid obtained from crossing two species *E. talpinus* (2n = 54) and *E. tancrei* (2n = 34). The scheme of partial karyotype shows three Rb metacentrics (1.2, 3.4, and 5.6) and six acrocentrics (1–6) like the intraspecific hybrid (D). *Sphere:* The main difference in the chromosome synapsis in SC trivalents is that the centromeric regions of the metacentrics are very stretched. The centromeric regions of acrocentrics are also stretched in some cases. Centromere stretching is associated with the difference in chromosome territories of two parent species. SC trivalents with stretched centromeres are presented on the right. Associations of sex chromosomes with SC trivalents (yellow stars) are often observed in both hybrids. Chromatin dense bodies (ChBs) of XX bivalents are a white ball on one of the X axes.

### Concluding remarks

Hybridization manifests a wide range of selective mechanisms, including decreased fertility or even sterility, due to defective synapsis and recombination (Miklos, 1974, de Boer, 1986, Davisson and Akeson, 1993), meiotic silencing failure (Homolka et al., 2007, Burgoyne et al., 2009), unbalanced chromosome segregation (Eaker et al., 2001), and alteration of nuclear architecture (Garagna et al., 2001). Modification of the nuclear architecture can be demonstrated by comparing native (parental) and experimental hybrids. The 34-chromosome mole voles differ in the positions of the chromosomal territories compared with the native 54-chromosome form due to 10 Rb pairs. To assess the differences in the location of chromosome territories, we performed experimental hybridization of 54- and 34-chromosome mole voles. We obtained two types of heterozygous mole voles with the same diploid number and identical chromosomal combinations. These two models are different in the manifestation of their fertility: intraspecific hybrids have slightly reduced fertility while interspecific hybrids are sterile. Forming the same number of SC trivalents, germ cells of the heterozygotes showed a different behavioral spectrum of chromosome combinations. In both hybrids, SC trivalents formed SC chains. However, in intraspecific hybrids, SC trivalents were able to leave such associations and form closed configurations, while in interspecific hybrids, only a few (usually 1–2) trivalents could complete the synapsis. The remaining SC trivalents could not dissociate from the SC chains, probably due to the peculiarities of the organization of centromeric regions in SC trivalents. The degree of nuclear architecture reorganization in the two hybrids was different, although the chromosome combinations were identical. These results suggest that alterations of nuclear architecture depend not only on the chromosome composition but also on other genetic or/and epigenetic factors ones.

Given that the behavior of the centromeric regions of SC trivalents was different in the two types of mole vole hybrids, it was logical to assume that this specificity could at least powerfully contribute to reproductive breakdown or be considered a cause of the sterility of interspecific hybrids. Because centromeres can be associated with patterns of gene expression and chromatin modifications (Zhao et al., 2016), physically stretched centromeric regions (as an indicator of nuclear architecture reorganization) could indirectly modify or destabilize these processes. Thus, we hypothesize that the shift in the intranuclear organization and the centromere stretching could change genetically significant regions of the genome through chromatin reformatting or altering the patterns of gene expression, which, in turn, could provoke the activation of meiotic checkpoints.

We suppose that pachytene irregularities and decreased recombination could synergistically, along with centromere instability and reorganization of intranuclear contents, contribute to the complete sterility of interspecific mole vole hybrids. In intraspecific hybrids, synaptic irregularities and a reduced crossover number could reduce fertility, but only in the F1 generation; future generations demonstrated ordinary fertility (Lyapunova and Yakimenko, 1985). The role of genetic factors in the development of sterility cannot be ruled out, because “hybrid sterility genes directly or indirectly modulate the sensitivity of synapsis to the sequence divergence between heterospecific chromosomes, either enhancing or suppressing it” (Bhattacharyya et al., 2013).

Summarizing our findings and previous studies, we conclude that a unique group of subterranean rodents, mole voles from the genus *Ellobius*, exhibits a wide range of interesting chromosomal phenomena, such as different systems of sex chromosomes, heterology of two isomorphic male X chromosomes, expressed in asynchronous chromatin inactivation at prophase I, multiple Rb translocations and monobrachial homologous metacentrics, neocentromere formation, stretched centromeres, acrocentric interactions leading to the formation of dicentric chromosomes and new Rb chromosome emergence. Each of the events could be considered as a driver or trigger of the karyotypic evolution of the rodents.

## Materials and Methods

### Animals

All Ellobius males were obtained from a mole vole collection of the IDB RAS. For meiotic studies, the parental Ellobius forms: three *E. talpinus*, two *E. tancrei* (2n=54), five *E. tancrei* (2n=34) and two groups of F1 hybrids: three intraspecific hybrids *E. tancrei* X *E. tancrei*, five interspecific hybrids *E. talpinus* X *E. tancrei* were used in this work.

The fertility of parental forms and hybrids was determined based on relevant data: 66 pups and 23 litters in *E. talpinus*, 168 pups and 71 litters in *E. tancrei* (2n=54), 197 pups and 82 litters in *E. tancrei* (2n=34), 41 pups and 20 litters in the intraspecific hybrid F1 *E. tancrei* X *E. tancrei*, 93 pups and 24 litters in the interspecific hybrid F1 *E. talpinus* X *E. tancrei*.

### Ethics statement

Experimental procedures were performed in strict accordance with the international, national, and institutional guidelines for animal care. Studies were approved by the Ethics Committee for Animal Research of the Vavilov Institute of General Genetics RAS (order 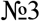 of November 10, 2016) and the Koltzov Institute of Developmental Biology RAS (Annual permissions, 2013-2017).

### Spermatocytes suspensions

The removed testes exempted from tunics, large blood vessels and fat. A piece of testis tissue was placed on cavity slide in 1 ml of Eagle’s medium (without glutamine), 37°C (Paneco, Russia). The testis tissue was crushed with razor blades, then the cell suspension was homogenized using an automatic pipette for 10-15 minutes. The degree of homogenization of the suspension and its cell composition was controlled under a light microscope.

### Spermatocytes spreads

Spermatocyte spreads were obtained in accordance the procedure of Kolomiets et al. (1986, 2010). 5 μl of testis cell suspension in Eagle’s medium was applied to the surface of the convex meniscus of a drop (20 μl) of 0.2 M sucrose solution (hypophase). The cells swelled up and then spread over the surface of the hypophase. After 1 min, a slide with a polylysine or coated with an electronically transparent chemically resistant film is touched to the surface of the hypophase (a 0.5% solution of the fragments of Falcon dish in chloroform). Cells adhering to the film surface were fixed with a 4% solution of paraformaldehyde, pH 8.0-8.4, washed with a 0.4% solution of Photoflo (Kodak), pH 8.0, and dried in air.

### Spermatocytes squashes

As emphasized by Berrios et al., 2010, squashes allow to preserve chromatin condensation and organization (3D space) of nuclei. Spermatocyte squashes were prepared in accordance the procedure of Page et al. (1998) (personal training from Jesus Page and Ana Gil-Fernandes in 2017). Removed testes were fixed in 2% formaldehyde in PBS with 0.05% Triton X-100 during 10 min. Then, several pieces of the seminiferous tubules were placed on a slide and squashed by exerting pressure on the coverslip. The slides were immersed in liquid nitrogen and the coverslips were removed with a knife. The slides were washed in PBS for 15 minutes and were immunostained.

### Antibodies and immunocytochemistry

The slides were washed in PBS. Spreads and squashes were blocked with HB (holding buffer: PBS, 0.3% BSA, 0.005% Triton X-100). The slides were incubated overnight at 4°C with primary antibodies: rabbit polyclonal antibodies SCP1 (Abcam, Cambridge, UK), rabbit polyclonal antibodies SCP3 (Abcam, Cambridge, UK) both diluted to a concentration of 1:500 in ADB (Antibody Dilution Buffer: PBS, 3% BSA, 0.05% Triton X-100), mouse antibodies MLH1 (1:50, Abcam, Cambridge, UK), mouse anti-phospho-histone H2AX (1:1000, Abcam, Cambridge, UK), human anti-centromere antibodies CREST (Fitzgerald Industries International, Massachusetts, USA), ACA (Antibody Incorporated, California, USA) or monoclonal CENPA (Abcam, Cambridge, UK), all diluted to a concentration from 1:200 to 1:400 in ADB. The slides were washed in PBS and incubated with goat anti-rabbit Alexa Fluore 488 conjugated antibodies (1:500, Abcam, Cambridge, UK), goat anti-human Alexa Fluore 546 conjugated antibodies (1:200 – 1:400) and goat anti-human Alexa Fluore 546 conjugated antibodies (1:500) at 37°C for two hours. The slides were washed with PBS, rinsed briefly with distilled water, dried and mounted in Vectashield with DAPI (Vector Laboratories). Some details of immunocytochemystry procedure see in Matveevsky et al., 2016, 2018. The slides were analyzed with an Axioimager D1 microscope CHROMA filter sets (Carl Zeiss, Jena, Germany) equipped with Axiocam HRm CCD camera (Carl Zeiss), and image-processing AxioVision Release 4.6.3. software (Carl Zeiss, Germany). Images were processed by Adobe Photoshop CS5 Extended

### AgNO3-staining and electron microscopy

The slides were stained with 50% AgNO3 solution in a humid chamber at 56°C for 3 hours. The slides were washed in four changes of distilled water and air-dried. The stained slides were observed in a light microscope, suitably spread cells were selected, and plastic (Falcon film) circles were cut out with a diamond tap and transferred onto grids. The slides were examined under JEM 100B or JEM 1011 electron microscope.

### Histological analysis of testis sections

Testes were removed, fixed overnight in Bouin’s solution, and then stored in 70% ethanol until use. The fixed testes were dehydrated in a graded ethanol series, immersed sequentially in ethanol / xylene and xylene, and then embedded in paraffin. The testes were sectioned at a thickness of 6 mm and mounted on glass slides. The sections were deparaffinized and stained with hematoxylin and eosin.

### Statistical analysis

The statistical analysis of all data was performed using GraphPad Prism 8 software (San Diego, CA, USA). Mean values (M) and standard deviation (SD) were calculated by descriptive option. P-values reported in Table S1 were calculated by Mann–Whitney two-sided non-parametric test. Bar charts, ring and dots diagrams were created by graph options of the software.

## Funding

This research was supported by the research grant of the Russian Foundation for Basic Research No 20-34-70027 (SM, AT, AK) and VIGG RAS State Assignment Contract (SM, OK); and by the Russian Foundation for Basic Research No 20-04-00618 (IB) and IDB RAS State Assignment for Basic Research (IB). Each of two RFBR grants supported only those parts of the work for which each of the co-authors was contribute. The funders had no role in study design, data collection and analysis, decision to publish, or preparation of the manuscript.

## Competing interests

The authors have declared that no competing intersts exist.

## Data Availability Statement

All relevant data are within the manuscript and its supporting information file.

## Acknowledgments

We thank the Genetic Polymorphisms Core Facility of the VIGG RAS, Moscow, for the possibility to use their microscopes. We thank S. Kapustina and V. Tambovtseva from IDB RAS for their help in maintaining the mole vole collection. We are grateful to GN Davidovich and AG Bogdanov of the Electron Microscopy Laboratory of Biological Faculty of Moscow State University for the technical assistance.

## Author Contributions

Maintenance of mole vole collection, hybridization, fertility data and karyotyping of Ellobius species and hybrids were performed by IB. All meiotic studies and calculation of gonadosomatic indices were carried out by SM, AT, AK and OK. Conceptualization of all meiotic researches of mole voles and its performing were supervised by OK. The visualization and presentation of all data and images design were done by SM. The primary original draft of the manuscript, including Supplementary materials, was written by SM and OK. All the authors read, discussed, modified and approved the final text of the submitted manuscript and the supporting information. All authors have read and agreed to the published version of the manuscript.

## Supplementary materials

**Table S1.** Total meiotic data of Ellobius species and hybrids

|  |  | 1 | 2 | 3 | 4 | 5 | 6 | 7 | 8 |
| --- | --- | --- | --- | --- | --- | --- | --- | --- | --- |
|  | Mole voles | Total number of pachytene nuclei | Number of cells with moved XX to the periphery of the nucleus, M±SD, % (n nuclei) [Fig 2I] | Open XX bivalents, M±SD, % (n nuclei) [Fig 2J] | Number of nuclei with associations of XX and SC trivalents (in hybrids) or XX and autosomes, M±SD, % (n nuclei) [Fig 2K] | Number of nuclei without free closed SC trivalents (SC trivalents in the chains), M±SD, % (n nuclei) [Fig 3E] | Number of nuclei with stretched centromeres, M±SD, % (n nuclei) [Fig 3I] | Number of nuclei with chromosome #7 detected, M±SD, % (n nuclei) [Fig 3L] | Average number of MLH1 foci per nucleus, M±SD (n nuclei) [Fig 5F] |
| A | *E. talpinus*, ♂, 2n=54, NF=56 (N=3) | 253 | 0.73±0.03, 72.59% (n=142) | 0.09±0.03, 9.85% (n=140) | 0 | – | 0 | always | 23.48±3.3 (n=101) |
| B | *E. tancrei*, ♂, 2n=54, NF=56 (N=2) | 153 | 0.72±0.07, 72.44% (n=132) | 0.11±0.07, 10.92% (n=132) | 0.06±0.04*, 5.9% (n=132) | – | 0 | always | 23.1±3.1 (n=99) |
| C | *E. tancrei*, ♂, 2n=34, NF=56 (N=5) | 241 | 0.76±0.09, 76.48% (n=98) | 0.09±0.02, 8.84% (n=98) | 0.02±0.03, 2.25% (n=98) | – | 0 | always | 22.8±3.7 (n=151) |
| D | Hybrid F1 *E. tancrei* X *E. tancrei*, ♂, 2n=44, NF=56 (N=3) | 168 | 0.50±0.11, 49.9% (n=108) | 0.53±0.11, 52.97% (n=114) | 0.50±0.08, 50.37% (n=115) | 0.09±0.02, 9.42% (n=99) | 0.16±0.06, 15.72% (n=96) | 0.94±0.02, 93.59% (n=78) | 15.4±4.4 (n=47) |
| E | Hybrid F1 *E. talpinus* X *E. tancrei*, ♂, 2n=44, NF=55 (N=5) | 198 | 0.49±0.13, 48.61% (n=172) | 0.55±0.15, 55.6% (n=113) | 0.45±0.18, 45.34% (n=98) | 0.78±0.12, 77.95% (n=102) | 0.69±0.11, 69.52% (n=105) | 0.91±0.09, 91.17% (n=102) | 14.8±4.7 (n=94) |
|  |  | Significant (P < 0.05) | A/D, A/E, B/D, B/E, C/D, C/E | A/D, A/E, B/D, B/E, C/D, C/E | B/D, B/E, C/D, C/E | D/E | D/E |  | A/E, B/E, C/E, A/D, B/D, C/D, D/E |
|  |  | Not significant | A/B, A/C, B/C, D/E | A/B, A/C, B/C, D/E | B/C, D/E |  |  | D/E | A/C, A/B, B/C |
|  |  | Comments |  |  | * We did not include data from the Matveevsky et al., 2015, in which this indicator is 0.17 ± 0.03. We used a different way of counting. |  |  |  |  |

**Fig S1.**
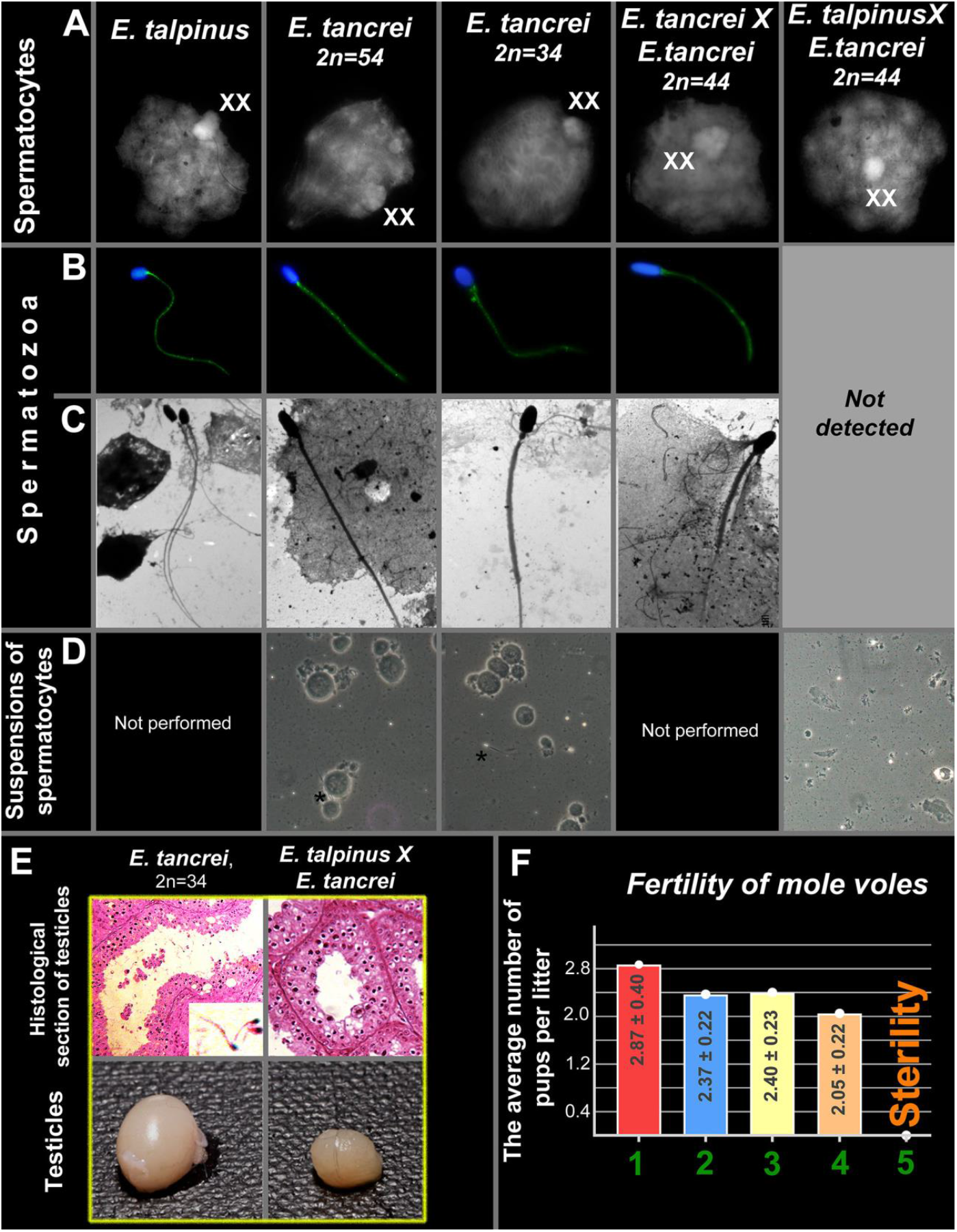
Fertility parameters of *Ellobius* species and hybrids. **(A)** DAPI-stained spermatocytes. **(B, C)** Spermatozoa: light (B) and electron (C) micrographs. Spermatozoa in B were detected by nonspecific immunostaining (SYCP3 antibody, green) and DAPI-staining (blue). **(D)** Suspension of spermatocytes. **(E)** Comparison of histological sections of testes and the size of testes in *Ellobius tancrei* (on the left) and the interspecific hybrid (on the right). **(F)** The average number of pups per litter in parents and hybrids: 1. *Ellobius talpinus;* 2. *E. tancrei* (2n = 54); 3. *E. tancrei* (2n = 34); 4. *E. tancrei* × *E. tancrei* hybrid; 5. *E. talpinus* × *E. tancrei* hybrid.

**Fig S2.**
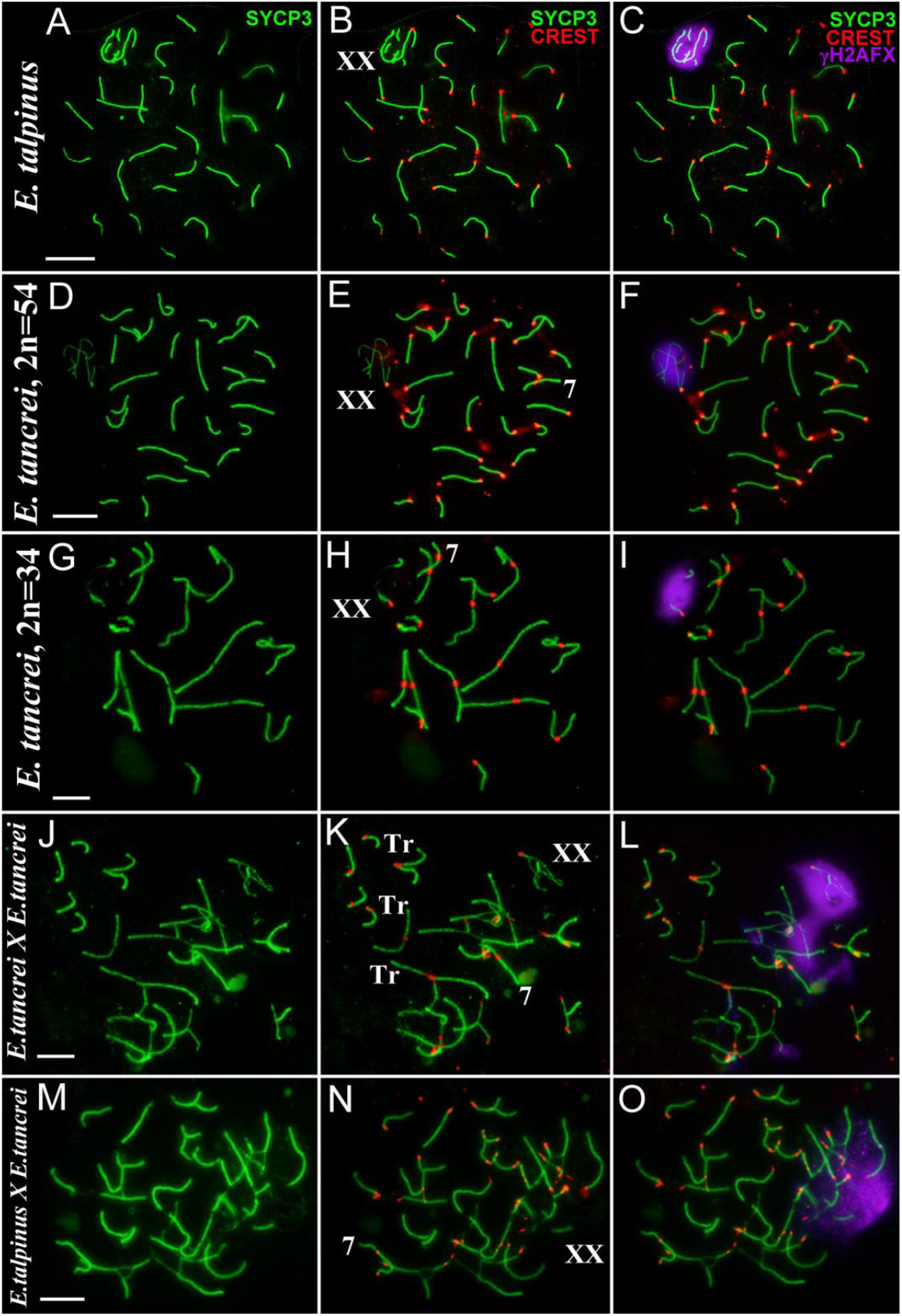
Pachytene spermatocytes of *Ellobius* species and hybrids. Synaptonemal complexes (SC)s were immunostained with antibodies against SYCP3 (green) and centromeres—with an antibody to kinetochores (CREST, red). An anti-γH2AFX (violet) antibody was used as a marker of chromatin inactivation. XXs were moved to the periphery of the nuclei and were covered by γH2AFX-cloud in parent species (**A–C**, **D–F**, and **G–I)**. The γH2AFX-cloud covered the XX bivalent and spread to some asynaptic segments in both 44-chromosome hybrids **(J–L** and **M–O)**. Scale bars represent 5 μm.

**Fig S3.**
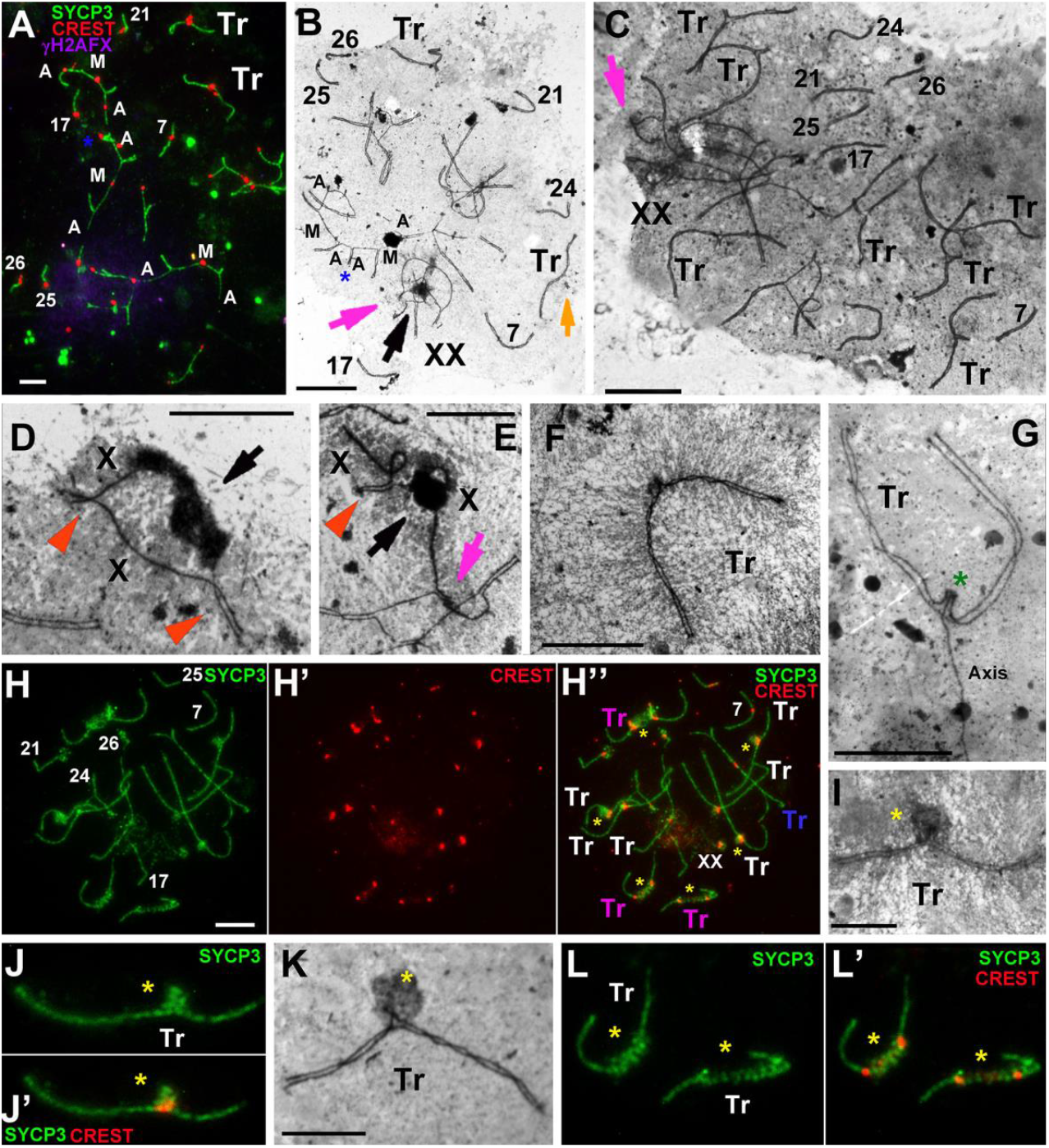
Synaptic behavior of chromosomes in intraspecific *E. tancrei* × *E. tancrei* hybrid. Light micrographs after immunostaining (A, H, J, and L). SCs were immunostained with antibodies against SYCP3 (green) and centromeres— with an antibody to kinetochores (CREST, red). An anti-γH2AFX (violet) antibody was used as a marker of chromatin inactivation. Electron micrographs after AgNO_3_-staining (B-G, I, and K). The numbers of autosomes (B, C, and H) correspond to the chromosome number of the karyotype (see Fig 1E). Orange arrowheads show synaptic sites of the sex (XX) bivalent. Black arrows show chromatin dense body (ChB) of the XX (B, D, and E). Tr indicates trivalent, M indicates metacentric, and A indicates acrocentric. **(A, B)** SC trivalents (A/M/A) chains joined together by short arms of nonhomologous acrocentrics at the early-to-mid pachytene stage (blue stars). XX bivalents have been associated with SC trivalents (Trs) or bivalents (see the violet signal in A and pink arrow in B). Both nuclei had two free Trs. The light orange arrow shows atypical prolonging of short arms of acrocentrics of the SC trivalent (B). **(C)** Seven free and closed SC Trs were formed at the late pachytene stage. XX bivalents were associated with SC chains (pink arrow). **(D)** Closed sex bivalent with two synaptic sites at the ends and a central unpaired region at the periphery of the nucleus. A long ChB was observed along one of the axes (black arrow). **(E)** One of the X chromosomes axes was in contact with the SC trivalent (pink arrow). **(F)** A closed SC trivalent with well-structured chromatin. **(G)** Triple synapsis in the region of short arms of a SC trivalent (green star). **(H**, **J**, **K**, and **L)** The only case of the nucleus with all 10 trivalents (H). SYCP3- and AgNO_3_-positive dense material was revealed in the region of short arms of some SC trivalents (yellow stars; white “Tr” in H; I–K). Some trivalents had a stretched metacentric, in the stretch segment of which SYCP3-positive dense material was also observed. Most nuclei had no such features in hybrid spermatocytes (L, L’). Scale bars represent 5 μm.

**Fig S4.**
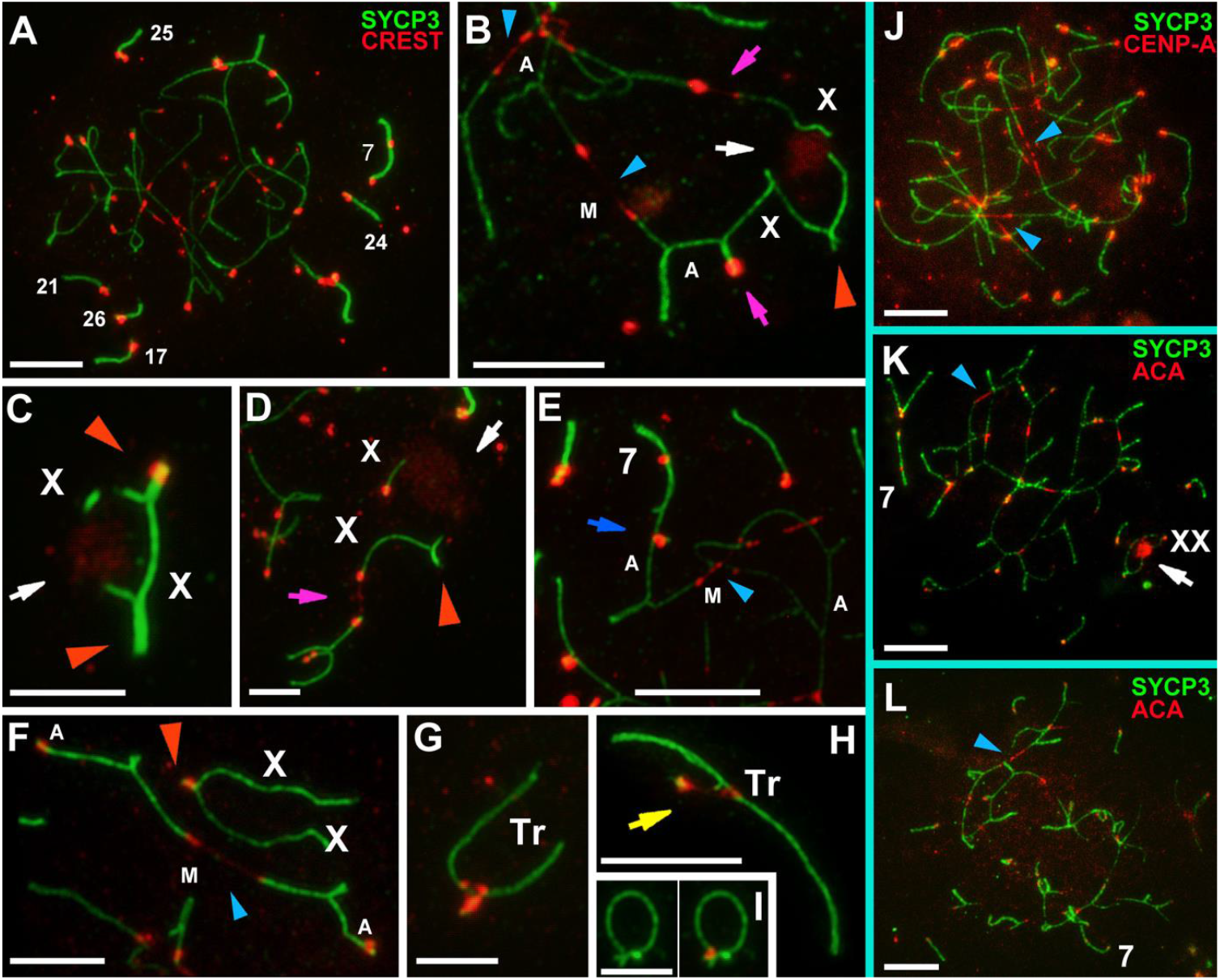
Synaptic behavior of chromosomes in interspecific *E. talpinus* × *E. tancrei* hybrid. SCs were immunostained with antibodies against SYCP3 (green) and centromeres—with antibodies to kinetochores (ACA, CREST, or CENP-A, red). The numbers of autosomes (A) correspond to the chromosome number of the karyotype (Fig 1D). Orange arrowheads show synaptic sites of the sex (XX) bivalent (B, C, D, and F). White arrows show chromatin dense body (ChB) of the XX (b, c, d, and k). Tr –indicates trivalent, M indicates metacentric, and A indicates acrocentric. **(A)** There were no free SC trivalents in the pachytene nucleus; all SC trivalents were in chains. The XX bivalent is not visible. **(B)** Double association of the XX bivalent with SC trivalents. An X univalent was associated with a SC trivalent by a thin CREST-positive line. **(C)** The XX bivalent with two synaptic sites and ChB. **(D)** An open XX bivalent with ChB. One of X univalent was associated with SC trivalent by a thin CREST-positive line. **(E)** The only case of the association of chromosome #7 with a SC trivalent. An acrocentric homolog of chromosome #7 involved in an association (blue arrow). **(F)** An open XX bivalent and SC trivalent with a stretched centromere. **(G)** A free closed SC trivalent. **(H)** The yellow arrow shows atypical lengthening of short arms of acrocentrics of the SC trivalent (H). **(I)** A ring-like univalent. **(J–L)** CENP-A-positive (J) and ACA-positive (K and L) centromeric region of metacentrics in some SC trivalents were stretched. There were no free SC trivalents. Scale bars represent 5 μm.

**Fig S5.**
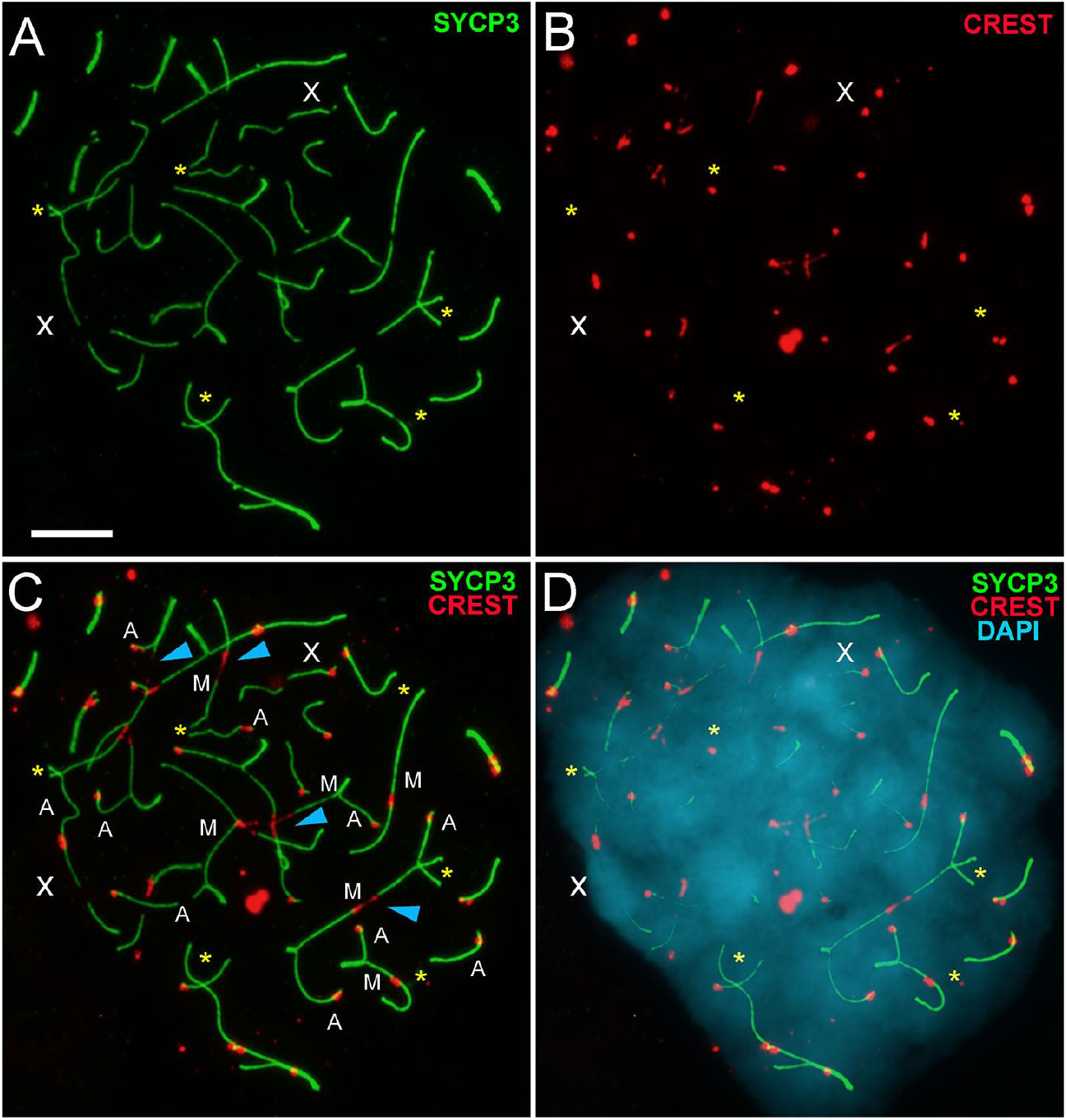
Synaptic behavior of chromosomes in interspecific *E. talpinus* × *E. tancrei* hybrid at zygotene nucleus. **(A-D)** SCs were immunostained with antibodies against SYCP3 (green) and centromeres—with an antibody to kinetochores (CREST, red). DAPI stained chromatin (blue). M indicates metacentric and A indicates acrocentric. Acrocentrics and metacentrics in SC trivalents adjusted together. Some acrocentrics and some arms of metacentrics had already entered synapsis, and some were just starting the synapsis process (yellow stars). Some metacentrics of SC trivalents had stretched centromeres (blue arrowheads). Sex chromosomes did not enter synapse and were located as X univalents. Their identification was possible based on length and a similar scenario of the behavior of axial elements, DAPI-positive signals, chromatin-dense body (ChB), and comparison with other micrographs (see Fig 2H). Scale bars represent 5 μm.

**Fig S6.**
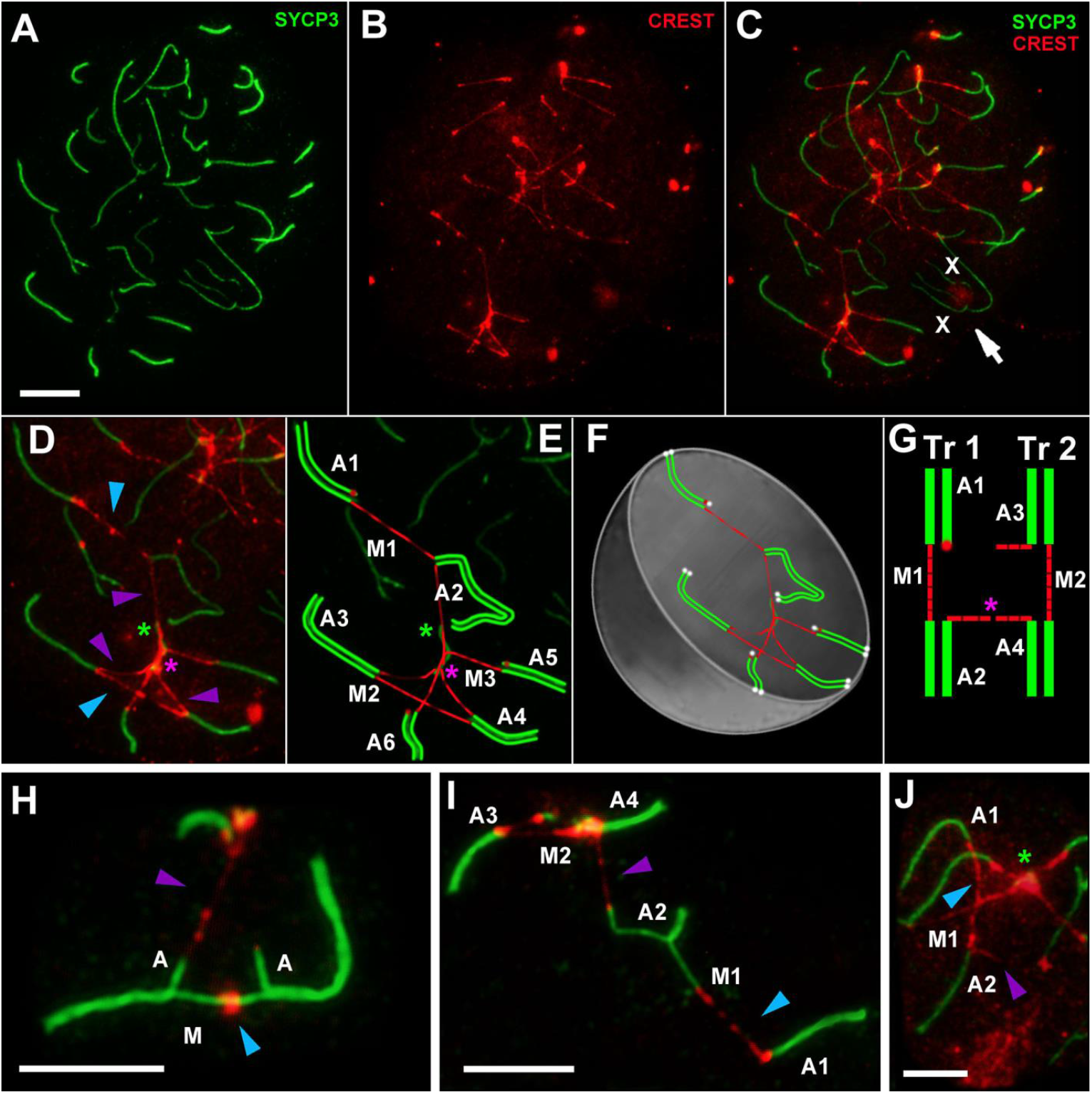
SC trivalents and stretched centromeres in interspecific *E. talpinus* × *E. tancrei* hybrid. SCs were immunostained with antibodies against SYCP3 (green) and centromeres—with an antibody to kinetochores (CREST, red). Blue arrowheads show centromeric regions of metacentrics in SC trivalents. Violet arrowheads show centromeric regions of acrocentric in SC trivalents. The white arrow shows a chromatin dense body (ChB) of the XX bivalent. Green stars show interlocking points. Tr indicates trivalent, M indicates metacentric, and A indicates acrocentric. **(A–C)** Pachytene nucleus: SYCP3-fragments (pseudobivalents) of SC trivalents were distant from each other (A). These fragments were connected by extensive stretched centromeric regions (B, C). **(D–G)** Part of the nucleus in A–C: Tree SC trivalents were linked by stretched centromeres of metacentrics and acrocentrics. SYCP3-segment of axial elements (green star) was an interlocking point. The scheme of the SC trivalent chains (E) and simulation of this chain in the nucleus (F) are presented. White dots at the chromosome ends mark the attachment points of the chromosomes to the nuclear envelope (F). Two SC trivalents A1/M1/A2 and A3/M2/A4 are connected by stretched centromeres of acrocentrics (pink stars) (D-G). **(H–J)** Examples of SC trivalents with stretched centromeres of metacentrics and acrocentrics. Scale bars represent 5 μm

**Fig S7.**
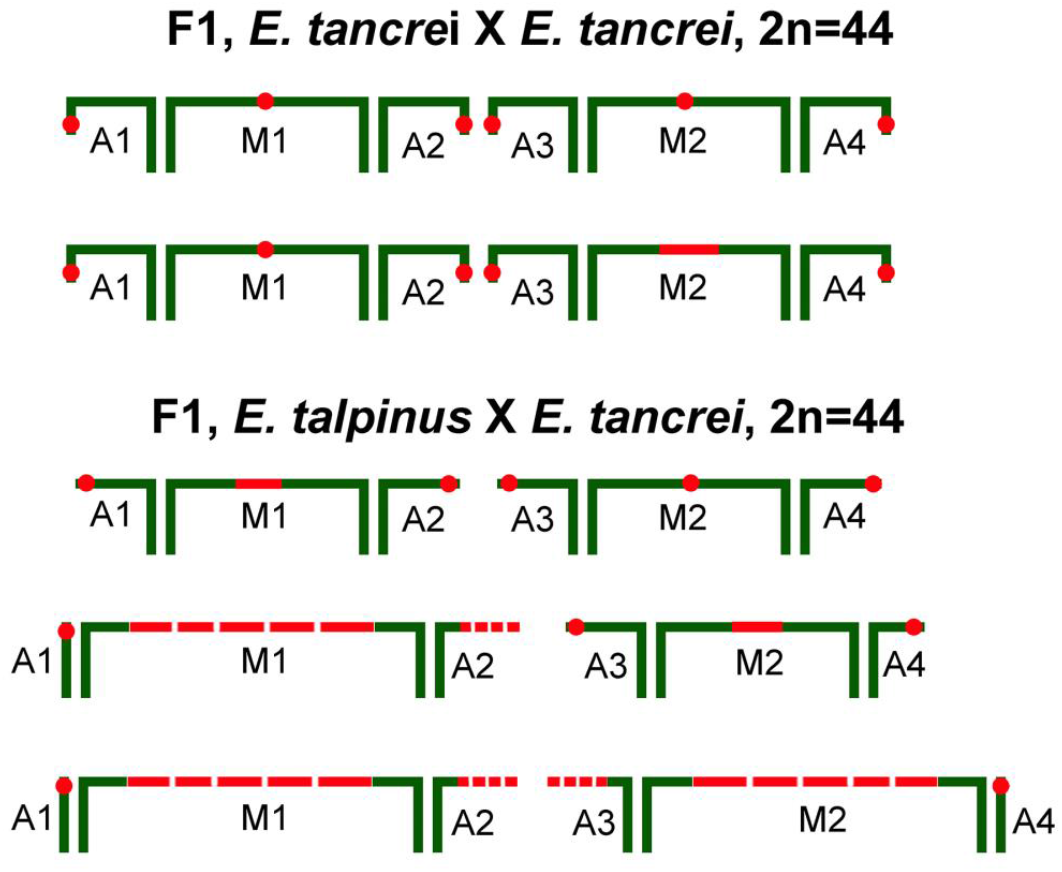
Types of SC trivalent chains in *Ellobius* hybrids. Red dashed inserts correspond to centromere stretching. Green lines correspond to the axial elements. See explanations in the text.

**Fig S8.**
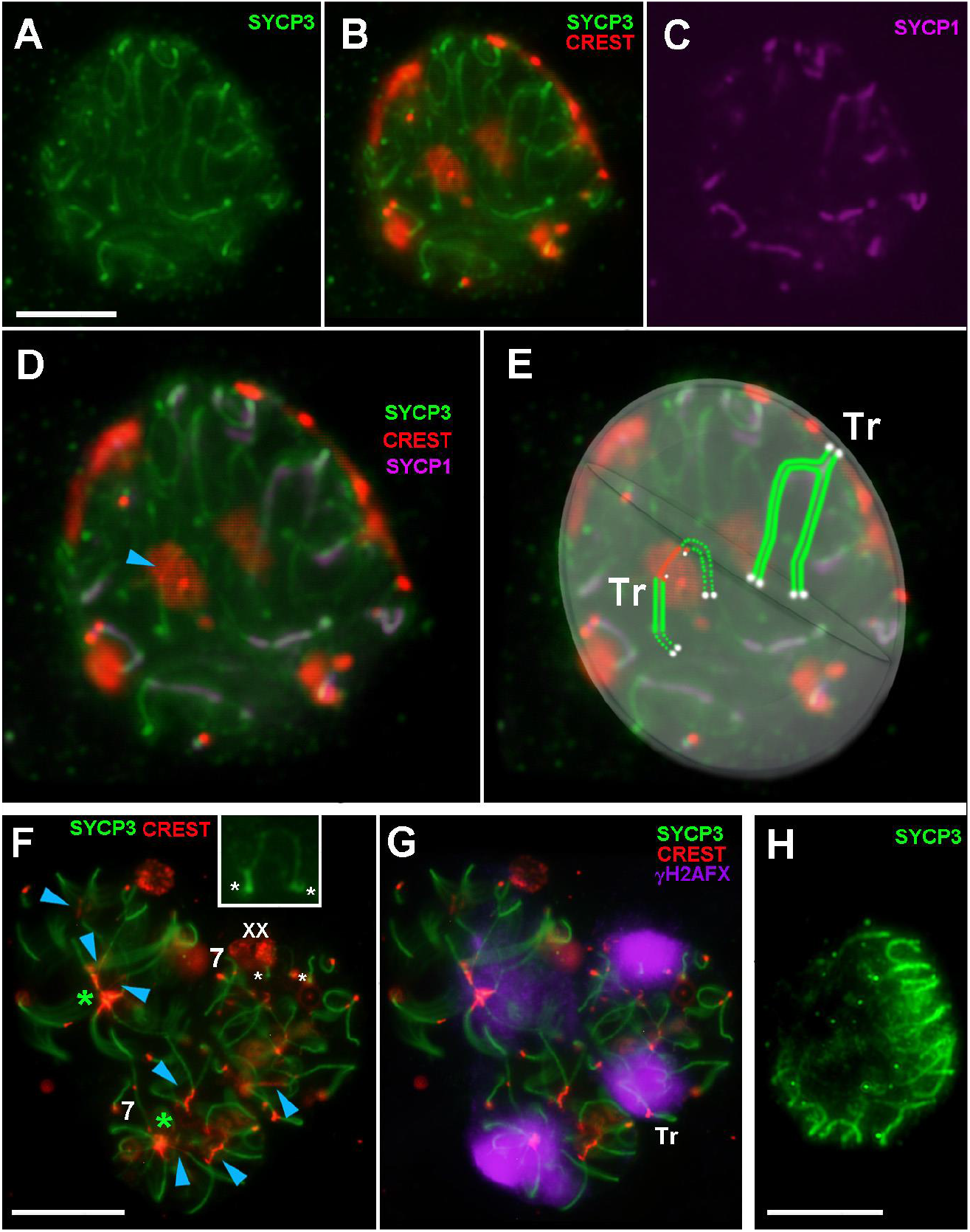
Spermatocyte squashes of interspecific *E. talpinus* × *E. tancrei* hybrid. Axial elements were identified using anti-SYCP3 antibodies (green), a central element using anti-SYCP1 (magenta), and an anti-CREST antibody for kinetochores (red). An anti-γH2AFX (violet) antibody was used as a marker of chromatin inactivation. DAPI stained chromatin (blue). Blue arrowheads show stretched centromeric regions in SC trivalents. One of the SC trivalent had a stretched centromere. **(A–E)** The short arms of the acrocentrics in the SC trivalents were attached to the nuclear envelope. The central element (SYCP1) was formed in the SC trivalent. White dots at the chromosome ends mark the attachment points of the chromosomes to the nuclear envelope (E). **(F**, **G)** Three squashes with some stretched centromeres of a SC trivalent. One of the squashes had a closed XX bivalent covered by a γH2AFX cloud. White stars show synaptic sites of the XX bivalent (enlarged in inset). Some open SC trivalents were also covered by a γH2AFX cloud (G). Some SC trivalents had a centromeric link in interlocking point (green stars) (F). **(H)** Zygotene-like stage: Chromosome–nuclear envelope interactions were visible. Scale bars represent 5 μm.

**Fig S9.**
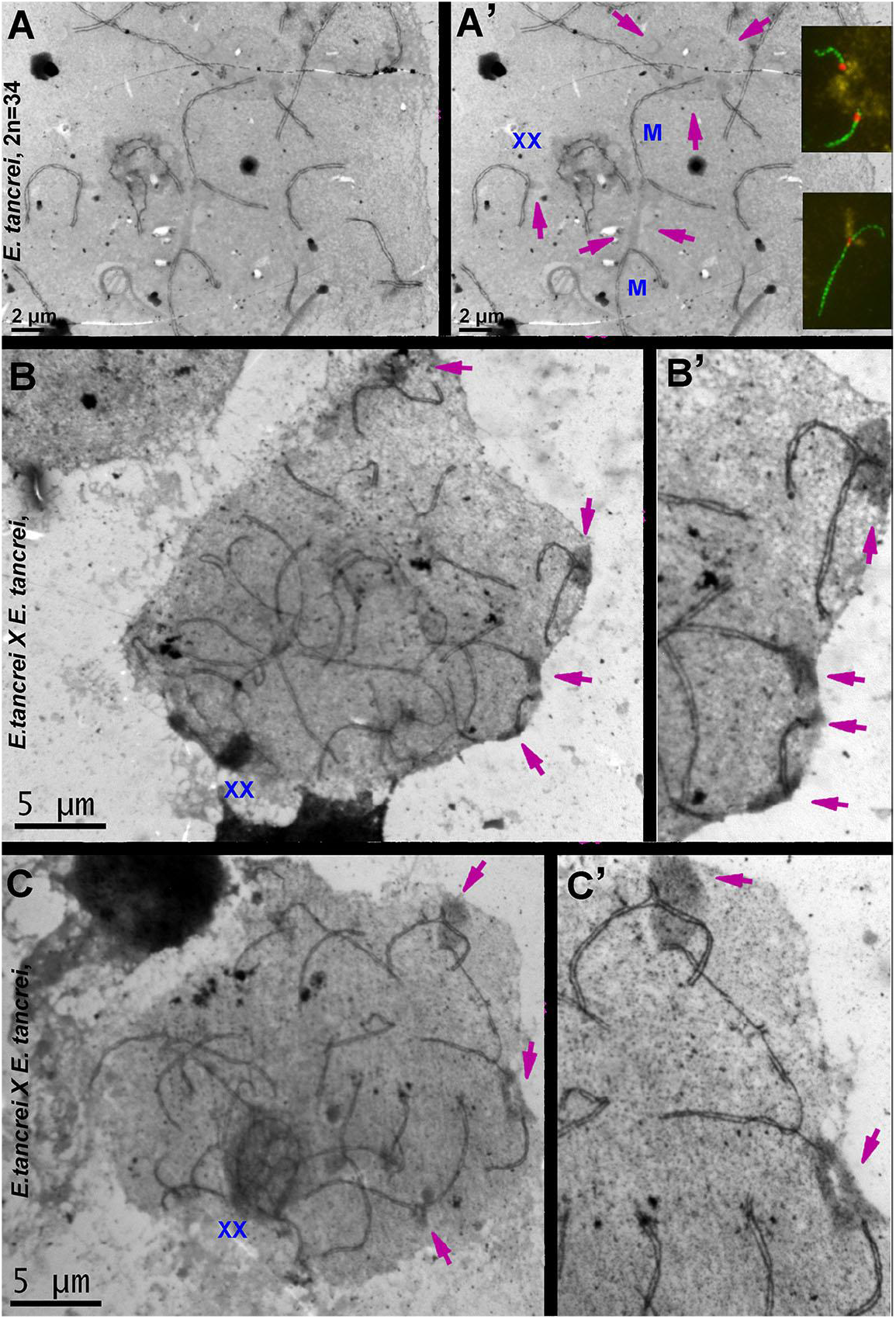
Heterochromatic links of chromosomes and nuclear envelope. Pink arrows show heterochromatin mass of chromosomes and SC trivalents. **(A**, **A’)** Part of the nucleus of *E. tancrei* (2n = 34) at the early pachytene stage. The linear heterochromatin link between two metacentrics (M) and cloud-like heterochromatic mass is visible. (Insets) SCs were immunostained by antibodies against SYCP3 (green), heterochromatin—by an H3K9me3 antibody (yellow)—and centromeres—by an antibody to kinetochores (CREST, red). H3K9me3 foci are cloud-like signals near the centromeres of acrocentrics (top inset) and linear signal near the centromere of the submetacentric (bottom inset). **(B**, **C)** Pachytene nuclei of the *E. tancrei* × *E. tancrei* hybrid. Some closed SC trivalents have a heterochromatic spot in the short arms of acrocentrics linked with the nuclear envelope (B’, C’). Heterochromatic spots of some acrocentrics have links with the nuclear envelope too (B’, C’).

## Notes

### Competing Interest Statement

The authors have declared no competing interest.

